# Atlas of genetic effects in human microglia transcriptome across brain regions, aging and disease pathologies

**DOI:** 10.1101/2020.10.27.356113

**Authors:** Katia de Paiva Lopes, Gijsje J. L. Snijders, Jack Humphrey, Amanda Allan, Marjolein Sneeboer, Elisa Navarro, Brian M. Schilder, Ricardo A. Vialle, Madison Parks, Roy Missall, Welmoed van Zuiden, Frederieke Gigase, Raphael Kübler, Amber Berdenis van Berlekom, Chotima Böttcher, Josef Priller, René S. Kahn, Lot D. de Witte, Towfique Raj

## Abstract

Microglial cells have emerged as potential key players in brain aging and pathology. To capture the heterogeneity of microglia across ages and regions, and to understand how genetic risk for neurological and psychiatric brain disorders is related to microglial function, large transcriptome studies are essential. Here, we describe the transcriptome analysis of 255 primary human microglia samples isolated at autopsy from multiple brain regions of 100 human subjects. We performed systematic analyses to investigate various aspects of microglial heterogeneities, including brain region, age and sex. We mapped expression and splicing quantitative trait loci and showed that many neurological disease susceptibility loci are mediated through gene expression or splicing in microglia. Fine-mapping of these loci nominated candidate causal variants that are within microglia-specific enhancers, including novel associations with microglia expression of *USP6NL* for Alzheimer’s disease, and *P2RY12* for Parkinson’s disease. In summary, we have built the most comprehensive catalog to date of genetic effects on the microglia transcriptome and propose molecular mechanisms of action of candidate functional variants in several neurological and psychiatric diseases.

## Introduction

Microglia, the myeloid immune cells of the brain, are a cell type of compelling interest in the pathogenesis of several brain disorders^1–3^. Microglia play critical roles in inflammatory responses, regulation of brain homeostasis, neurodevelopment, and neurogenesis. Microglia are highly dynamic cells that are strongly influenced by different environmental signals which result in distinct phenotypes and functions across brain regions^4–9^. In addition, microglial functions vary across different ages, disease pathologies, and between sexes^8,10–15^. For decades, changes in microglial density, morphology, and transcriptional state have been observed in postmortem brain tissue of patients with neurological and psychiatric disorders^16–20^. This was initially suggested to reflect a response of the immune system to underlying disease processes. However, recent evidence from genome-wide association studies (GWAS) and other follow-up analyses has suggested that a proportion of the genetic risk of neurological and psychiatric diseases acts through myeloid cells^21–24^. As the myeloid cells of the nervous system, microglia may therefore play a causal role in disease. However, due to the current scarcity of human microglia data, only a small number of genetic variants have been localized specifically in microglia, such as *BIN1* in Alzheimer’s Disease (AD)^25,26^.

There is a critical need to associate genetic risk variants to changes in the microglial transcriptome through quantitative trait loci (QTLs) to better understand the role of microglia in disease processes in the brain and to identify microglia-related targets for treatment in the long-term. So far, investigations of genetically-driven changes in gene expression and splicing in microglia have been limited by the availability of material from the number of subjects required to perform well-powered genomic analyses. Recently, Young *et al*. constructed expression QTLs (eQTLs) in primary human microglia (*n* = 93), and detected 401 eQTLs, some of which colocalized with AD loci, including *BIN1*^*27*^. However, the microglial transcriptome is highly heterogeneous compared to other cell types^28^, so larger sample sizes are needed to detect statistically significant eQTLs. In addition, it is becoming increasingly clear that genetic risk can also be mediated through mRNA splicing^29,30^. For instance, the *CD33* locus in AD influences *CD33* splicing, resulting in isoforms with different biological and likely pathological functions^30^.

In the present study, we describe the Microglia Genomic Atlas (MiGA), a genetic and transcriptomic resource comprised of 255 primary human microglia samples isolated *ex vivo* from four different brain regions of 100 human subjects with neurodegenerative, neurological, or neuropsychiatric disorders, as well as unaffected controls (**Figure 1**). We performed systematic analyses to investigate sources of microglial heterogeneity, including brain region, age, and sex. We further performed expression and splicing QTL analyses in each region and performed a meta-analysis across the four regions to increase our discovery power. We then performed colocalization and used fine-mapping and microglia-specific epigenomic data to prioritize genes and variants that influence neurological disease susceptibility through gene expression and splicing in microglia. With this approach, we have built the most comprehensive resource to date of *cis*-genetic effects on the microglial transcriptome and propose underlying molecular mechanisms of potentially causal functional variants in several brain disorders.

**Figure 1.**
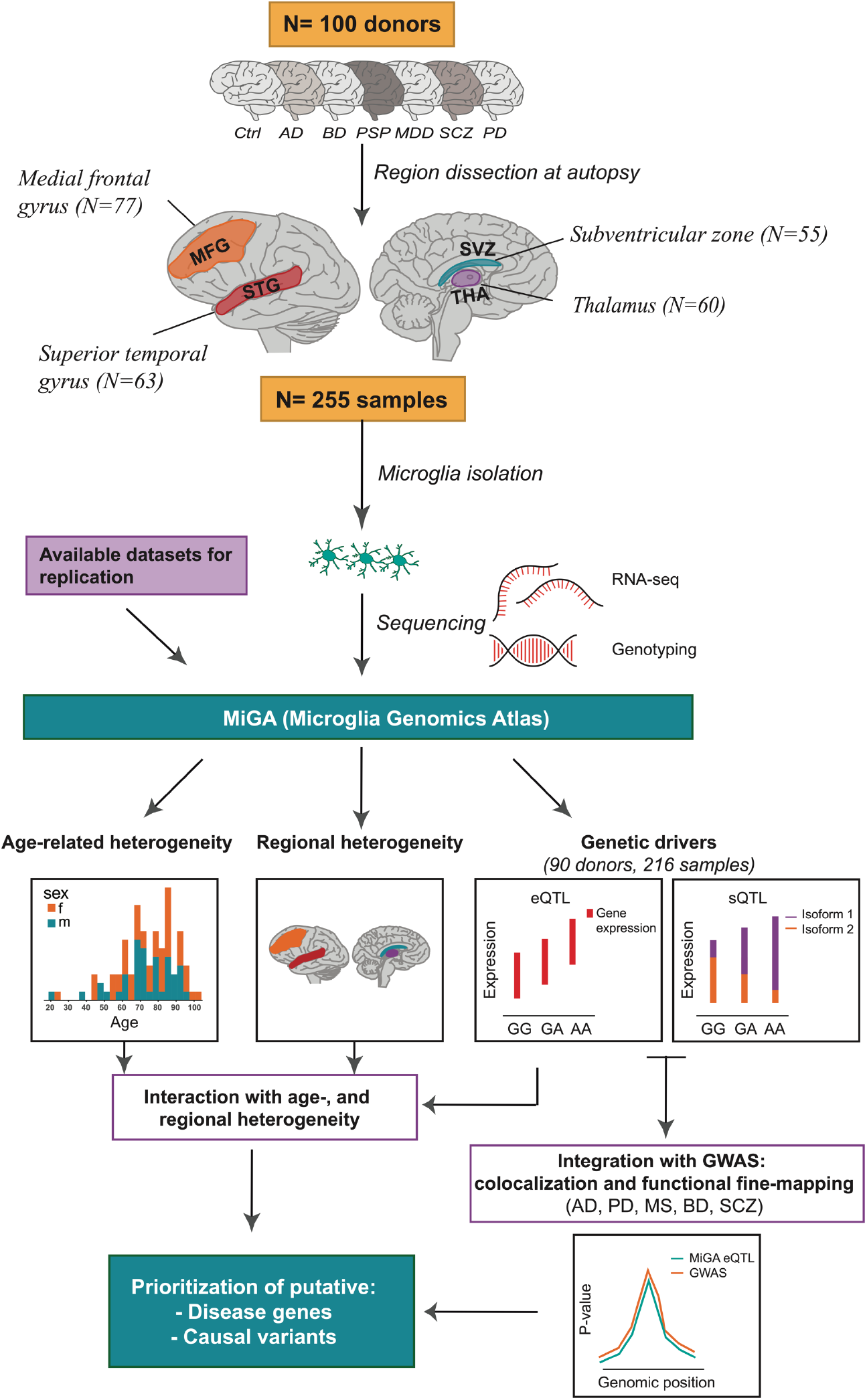
Overview of the Microglia Genomic Atlas (MiGA). Primary human CD11b+ microglia were isolated at autopsy from 100 donors with neurological and psychiatric diseases, as well as unaffected subjects (controls) generating a total of 255 samples from four brain regions: medial frontal gyrus (MFG), superior temporal gyrus (STG), subventricular zone (SVZ) and thalamus (THA). Samples were isolated from two brain banks: the Netherlands Brain Bank (NBB) and the Neuropathology Brain Bank and Research CoRE at Mount Sinai Hospital. RNA was isolated and sequenced. Genome-wide genotyping was performed using DNA isolated from all donors. The following analysis were performed with the MiGA dataset: (i) age-related analysis; (ii) regional heterogeneity analysis by looking at the differentially expressed genes between the brain regions; (iii) expression quantitative trait loci (eQTL) analysis; (iv) splicing quantitative trait loci (sQTL) analysis; (v) colocalization and functional fine-mapping integrating the eQTL results with the most recent GWAS from five diseases: Alzheimer’s disease (AD), Parkinson’s disease (PD), Multiple sclerosis (MS), Bipolar disorder (BD) and Schizophrenia (SCZ).

## Results

We collected 314 human microglial samples from 115 donors, of which 46 donors were non-neurological disease controls (**Figure S1**). Microglia were isolated from fresh post-mortem tissue using CD11b-beads^31– 33^. Mass cytometry (CyTOF) proteomic analysis showed an average of 95% of cells being positive for the microglia-specific marker P2Y12 (**Figure S2A**). The microglial samples were all subjected to RNA-sequencing using a low input library preparation. After rigorous quality control, 255 microglial samples from 100 different donors and four different regions were included (**Figure S1; Figure S3; Table S1)**. The four regions comprise two cortical regions: the medial frontal gyrus (MFG) and superior temporal gyrus (STG); and two subcortical regions: the thalamus (THA) and subventricular zone (SVZ). All microglial samples expressed known microglia-specific genes at high levels, with marker genes for neurons, astrocytes, and oligodendrocytes being lowly or not expressed (**Figure S2B**).

### Regional and aging heterogeneity in the microglia transcriptome

We explored the variation of a wide range of biological factors in driving the human microglia transcriptome before and after controlling for technical confounders (**Figure 2A; Figure S4-S5**). Using principal component analysis (PCA), we observed no clear separation by any factor after regressing out the technical factors, with the exception of age, which stands out clearly in PC2 (**Figure S6**). Sex explained little variance (**Figure 2A**), and we observed no differentially expressed genes between males and females (**Table S2**). Using a linear mixed model to estimate the variance explained for a set of factors per gene^34^, we found that donor identity explained the most variance per gene (mean 13.5%) (**Figure 2A)**. Brain region explained comparatively little variance overall (mean 2.95%), but we identified a subset of genes that were strongly variable between regions. We performed pairwise comparisons of differential gene expression between each pair of regions, accounting for shared donors in a linear mixed model^35^ (**Figure 2B-C, Table S3-S8**). The largest number of differentially expressed genes (FDR < 0.05; |log_2_ fold change| > 1) were between the SVZ and the two cortical regions (609 in MFG, 909 in STG), whereas comparing STG to MFG found the fewest (6 genes), suggesting microglia in the two cortical regions to be similar. We compared our findings in a published dataset of white and grey matter microglia^5^ and found significant overlaps with our MFG vs SVZ comparison (upregulated OR = 18.4; *P* < 1 x 10^-16^, downregulated OR = 4.83; *P* = 9 x 10^-6^; Fisher’s exact test; **Figure 2D**).

**Figure 2.**
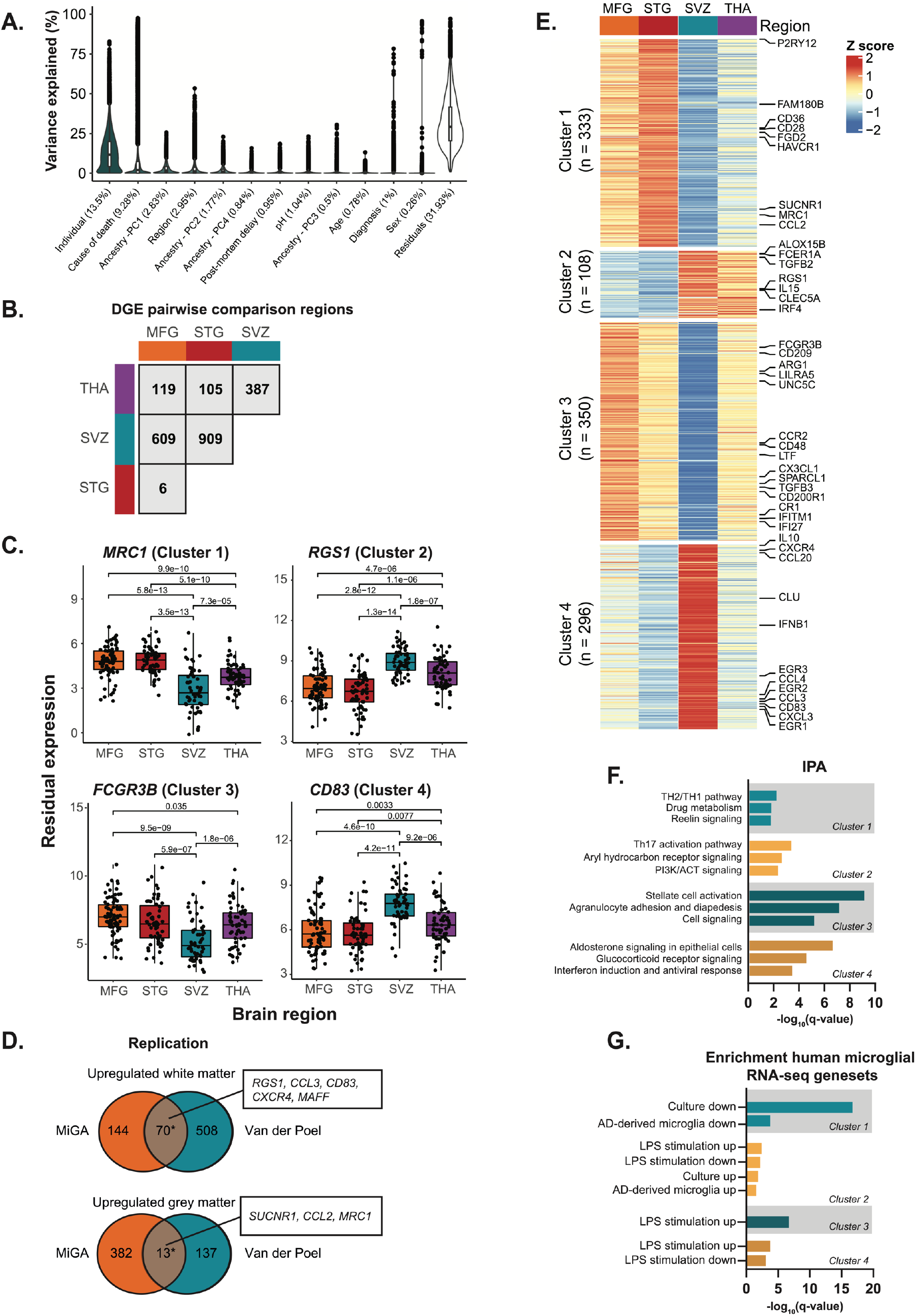
Regional heterogeneity analysis. A) Distributions of variance explained per gene for the non-technical factors. Mean variance explained by each factor is in brackets. B) Number of differentially expressed genes by pairwise region comparison (FDR < 0.05 and |logFC|>1). C) Examples of differentially expressed genes. The x-axis shows each brain region and the y-axis shows the residual expression after regressing out covariates. The *P*-values are from the linear regression using the Dream package after adjusting by FDR. D) Replication analysis with an independent dataset of microglia samples isolated from white (n = 5) and grey matter (n = 11) of controls^5^. Asterisk indicates significant enrichment by Fisher’s exact test (*P*-value < 0.05). Selected genes in overlap are highlighted. E) Heatmap of K-means clustering of 1,087 differentially expressed genes from the pairwise comparison. K-means was performed on z-scored values of the median per region of *voom* transformed expression. The colors represent row scaled z-score levels: red and blue indicates high and low relative region expression, respectively. F) Functional Enrichment Analysis of each K-means cluster from E using the software Ingenuity Pathway Analysis (IPA). The x-axis shows the significance, -log_10_(q-value) and y-axis shows only the significant terms (FDR q-value < 0.05). G) Functional Enrichment Analysis with curated human microglia RNASeq gene sets^37,38^ using Fisher exact test at Bonferroni-adjusted *P* < 0.05. The x-axis shows the significance, -log_10_(q-value) and y-axis shows the curated human microglia RNASeq gene sets (q-value < 0.05).

We then performed k-means clustering^36^ of the genes found in the pairwise comparisons. Testing for different numbers of clusters, we identified k = 4 as the optimal clustering partitioning after minimizing the total within-cluster sum of squares (WSS). That allowed us to identify four distinct groups of genes with specific expression patterns across brain regions (**Figure 2E**). Cluster 1 contains genes that are upregulated in the cortical regions compared to subcortical regions, such as *P2RY12, CD36*, and *MRC1*. Cluster 2 contains genes that are downregulated in the cortex compared to the subcortex (e.g. *FCER1A, IL15)*. Cluster 3 contained genes specifically downregulated in the SVZ (e.g. *CX3CL1, CCR2*) and cluster 4 contained genes upregulated in SVZ (e.g. *IL10, CLU)*, compared to the other three regions. We found that genes implicated in inflammatory processes were highly expressed in cluster 2 (**Figure 2F**), whereas genes related to homeostatic functions of microglial cells were mainly present in cluster 1. Cluster 4 includes genes that are involved in biological functions related to hormonal signaling and interferon response (**Figure 2F**). We overlapped the region-specific genes with gene sets altered after stimulation with lipopolysaccharide (LPS) or interferon-gamma (IFNγ; generated in-house), following *in vitro* culture^37^, and in microglia derived from AD patient brains compared to controls^38^. Cluster 1 was enriched in genes that were downregulated in AD brain and in response to *in vitro* culture, whereas cluster 2 genes were enriched in genes that responded in the opposite direction. Clusters 2, 3, and 4 were enriched for different sets of LPS-responsive genes in both directions (**Figure 2G, Table S9**).

To assess the effect of aging on the microglial transcriptome, we fitted a linear mixed model accounting for shared donors across all four regions. We observed 1,693 genes (338 up, 1,355 down at FDR < 0.05) associated with the chronological age of subjects, including many microglial-specific genes (*P2RY12, TLR2, C2*) (**Figure 3A, Table S10**). Genes upregulated in aging were significantly enriched for a number of Gene Ontology (GO) biological processes including lipid metabolism, immune responses such as Natural Killer (NK) cell and interferon signaling, and phagosome formation (**Figure 3B**). The downregulated genes were significantly enriched for cell motility, polarity, IL-6 cytokine signaling (**Figure 3B**), and for genes also downregulated following *in vitro* culture ^37^ and in AD-derived microglia^38^ (**Figure 3C**).

**Figure 3.**
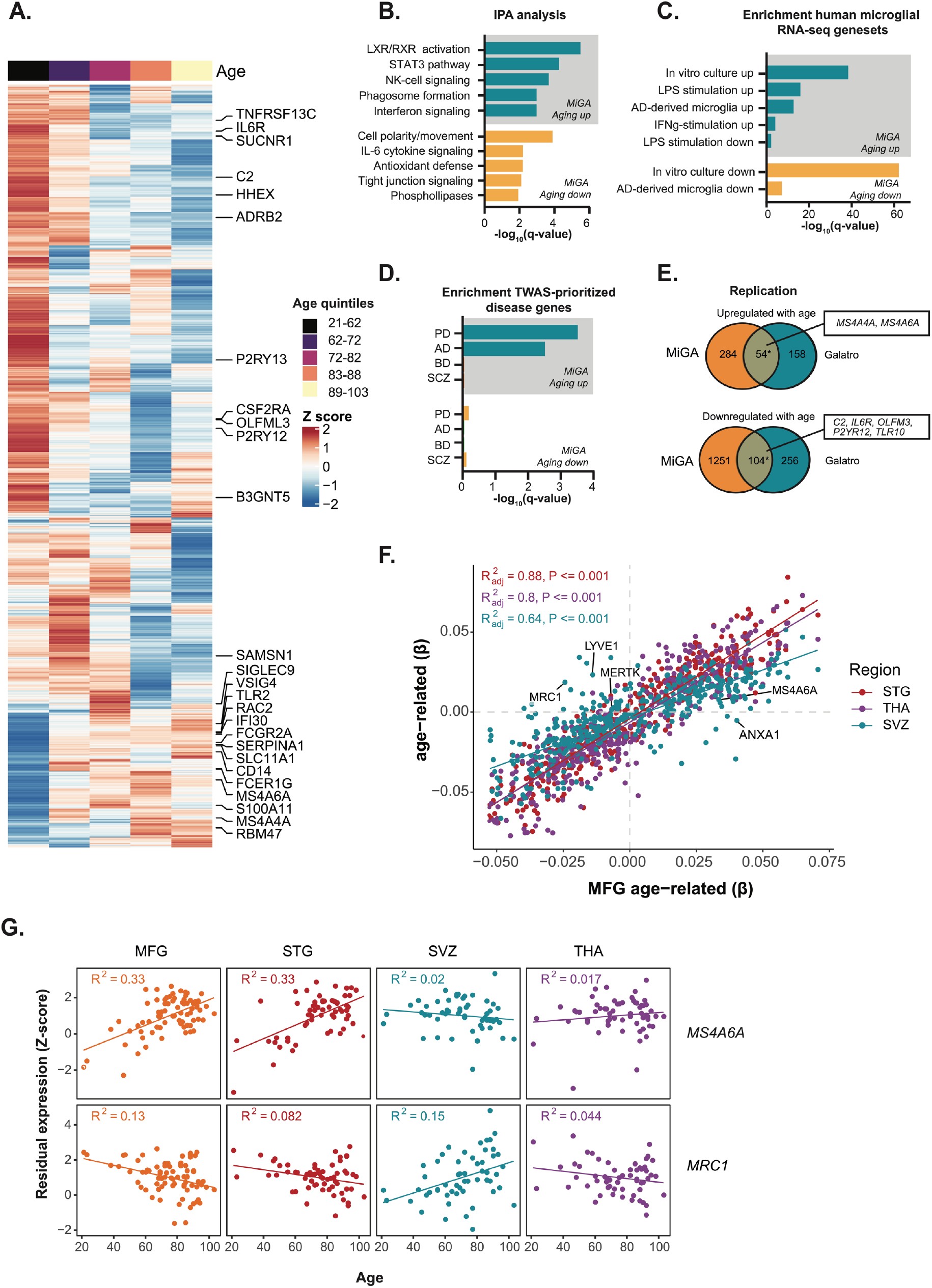
Age-related analysis. A) Heatmap showing the 1,693 genes associated with age from the linear regression using the Dream package (FDR < 0.05). Each row shows the z-scored median expression, averaged first by donor (across multiple regions) and then by quintiles with 20 donors each. Rows are ordered according to the Ward’s hierarchical clustering. The quintiles are ordered by age as indicated in the top bar, ranging from 21 to 103 years old. Colors in the heatmap represent row scaled z-score levels: red indicates high relative expression and blue indicates low relative expression. B) Pathway analysis of upregulated and downregulated age-related genes. The x-axis shows the significance, -log_10_(q-value) and y-axis shows only the significant terms (q-value < 0.05). C) Enrichment analysis with curated human microglia RNASeq gene sets^37,38^ using Fisher exact test at Bonferroni-adjusted *P* < 0.05. The x-axis shows the significance, -log_10_(q-value) and y-axis shows only the curated human microglia RNASeq gene sets (q-value < 0.05). D) Genes associated with age show overrepresentation for TWAS prioritized genes for Alzheimer’s (AD) or Parkinson’s disease (PD), but not for genes in Schizophrenia or Bipolar disorder. E) Replication analysis with an independent dataset of human microglia samples, 49 healthy controls with ages between 31 and 102 years old^15^. Asterisk indicates significant enrichment by Fisher’s exact test (*P*-value < 0.05). Selected overlapping genes are highlighted. F) Scatter plot showing the size correlation of the age-related genes by brain region. Only the significant genes (FDR < 0.05) in at least one region are shown. The x-axis shows the beta values in the MFG region and the y-axis shows the betas in other brain regions (STG, SVZ and THA). G) Scatter plot showing the z-scored residual expression for selected genes (*MRC1* and *MS4A6A*) by age and brain region. A distinct trajectory is observed in the SVZ compared to cortical regions.

We next used gene sets prioritized by transcriptome-wide association study (TWAS) in different diseases^39– 42^ (**Table S11**). The upregulated genes in chronological aging showed overrepresentation for genes in AD (4 out of 36 genes, e.g. *MS4A6A, FCER1G*, and *CR1*) or PD (7 out of 77 genes, e.g. *BST1, PTPN22, and TNFSF13)* GWAS loci, but not for genes in schizophrenia (SCZ) or bipolar disorder (BD) (**Figure 3D**). We replicated our findings using an external microglia aging dataset from the parietal cortex^15^ (upregulated genes OR = 23.4; *P* < 1 x 10^-16^, downregulated genes OR = 5.97; *P* < 1 x 10^-16^; Fisher’s exact test; **Figure 3E**), and from peripheral blood^43^ (**Figure S7**).

It is not known whether the impact of aging on the microglial transcriptome is uniform throughout the human brain. Although most genes showed concordant effect size and direction across regions (**Figure 3F**), 91 genes demonstrated age-region relationships after fitting an interaction model (FDR < 0.05) (**Figure S8A**). A set of them had increased (n = 16) or decreased (n = 11) expression during aging in MFG and STG, but not THA and SVZ (e.g. *MS4A6A*), while 25 genes (e.g. *MRC1, CD24*) changed specifically in SVZ and not in other regions (**Figure 3G; Figure S8B, Table S12**). Together, our results indicate that the microglial phenotype ages in a generally uniform manner across brain regions, with a distinct aging trajectory observed in a minority of genes.

### Genetic regulatory effects in microglia from multiple brain regions

We performed *cis*-eQTL and *cis*-sQTL analyses in primary human microglia from four different brain regions. After QC, 216 samples from 90 individuals of European ancestry were used for the analysis (**Figure S9**). In the region-specific analysis, we observed between 67 and 199 genes with a *cis*-eQTL (eGenes) and 253 to 426 genes with a *cis*-sQTL (sGenes) per region (FDR < 0.05; **Table S13, Figure S10**). As expected, *cis*-QTL discovery was highly correlated with the sample size for each region (Spearman’s *ρ* = 0.8 for eQTLs, and 1 for sQTLs). The low number of eQTLs detected in the region by region analysis is likely due to the small sample size per region and/or high donor-to-donor variation in microglia **(Figure 2A**). We therefore performed a meta-analysis across all four regions using the multivariate adaptive shrinkage (mashR)^44^ method to increase power and to assess shared QTLs between regions. In total, we identified 3,611 eGenes and 4,614 sGenes, at a local false sign rate (*lfsr*) < 0.05 in at least one region (**Figure 4A, Table S14-S15**).

**Figure 4.**
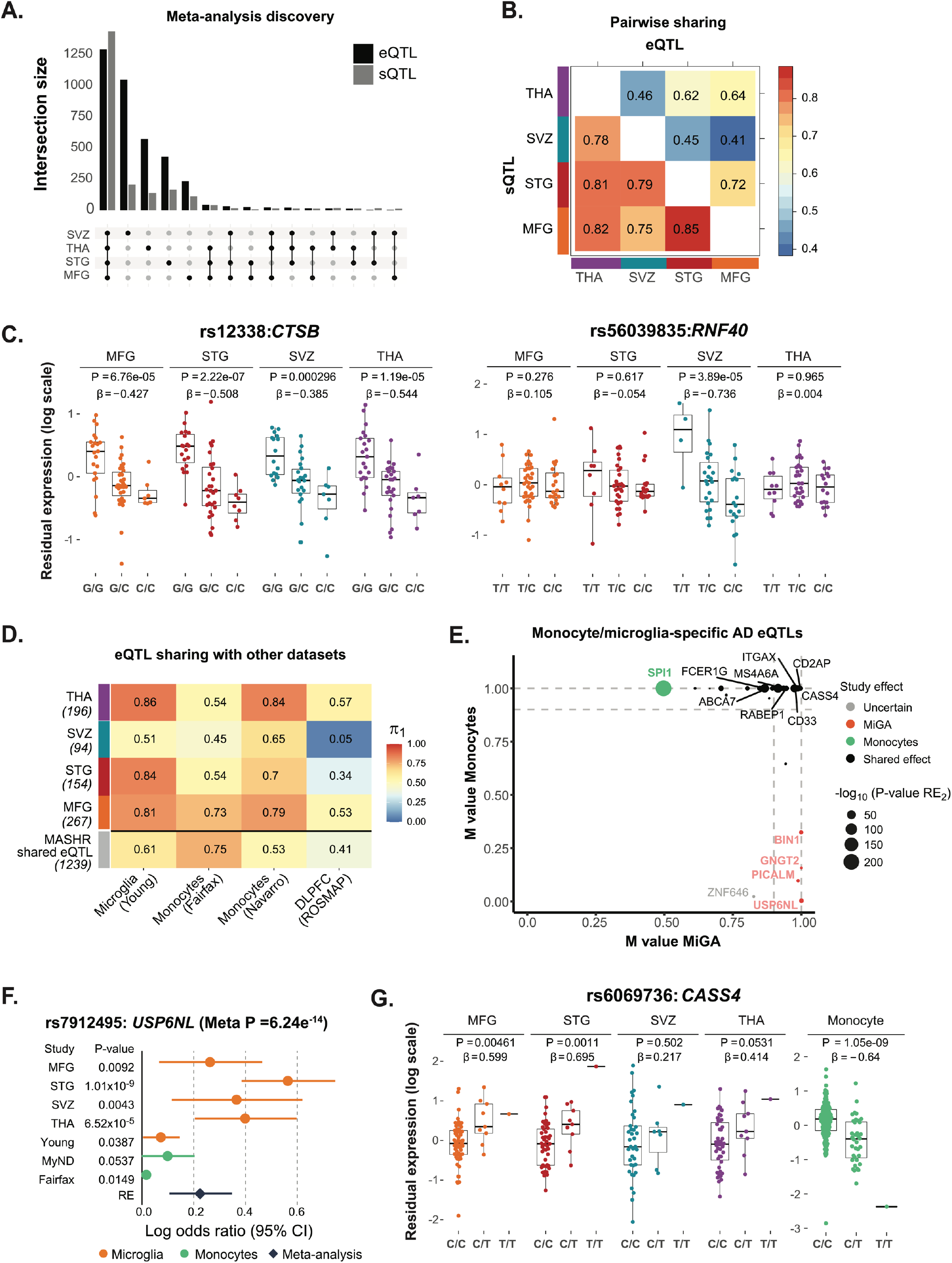
Genetic regulatory effects in microglia. A) Number of genes with a *cis*-eQTL (eGenes) and cis-sQTL (sGenes) at local false sign rate (*lfsr*) < 0.05, from a meta-analysis across all four brain regions using mashR. B) Pairwise shared eQTLs (upper triangle) and sQTLs (lower triangle) across the four brain regions. Numbers represent the proportion of significant effects (*lfsr* < 0.05) that are shared in magnitude (i.e. effect estimates that are in the same direction and within a factor of 2 in size). C) Examples of shared (*CTSB* gene, rs12338) and region-specific effect (*RNF40*, rs56039835). eQTL boxplots with residual gene expression (PEER adjusted) per individual stratified by genotype. The eQTL *P*-value and effect size are listed on top. D) Replication of MiGA eQTLs effects (region-by-region analysis q-value < 0.10 and mashR results at *lfsr* < 0.05) compared to four external eQTL datasets: microglia^45^, monocytes^46,47^, and dorsolateral prefrontal cortex (ROSMAP)^48^. The proportion of replication is measured by Storey’s π_1_. E) Meta-analysis results for colocalized eGenes in AD loci. The m-values represent the posterior probability that the effect exists in each study (i.e. MiGA for microglia, or Navarro and Fairfax for monocytes), as reported by the software METASOFT. Small m-value (< 0.1) means that the gene and the SNP do not have an effect in the study; large m-value (> 0.9) means that the gene and the SNP have an effect. Otherwise, the prediction is uncertain. The x-axis shows the maximum m-values among the four brain regions in MiGA, and y-axis shows maximum m-values for monocytes^46,47^. Colors indicate cell type: orange for the genes with strong effect in MiGA only, green for monocytes and black for the shared effects between microglia and monocytes. F) Example of microglia specific eQTL found only in MiGA. The gene *USP6NL* (rs7912495) has significant effect in all four brain regions from MiGA but does not have an effect in other datasets. G) Example of discordant eQTL effects for *CASS4* (rs6069736) between microglia and monocytes.

To investigate eQTL and sQTL sharing between microglia from different brain regions, we assessed the sharing of effects by magnitude (effect estimates that are in the same direction and are similar sizes, within a factor of 2). We observed a high degree of eQTL sharing between MFG and STG (72%), as expected, given that these two cortical regions have similar gene expression patterns (**Figure 4B**, upper triangle). Microglia from SVZ exhibited lower pairwise sharing of eQTLs with other regions, with the lowest sharing by magnitude observed between SVZ and MFG (41%), consistent with observed transcriptomic differences between these two regions. For sQTLs, we found overall higher regional sharing effects compared to eQTLs, but still following the same trends, with SVZ and MFG as the lowest (75%) and MFG and STG the highest (85%) pairwise sharing (**Figure 4B**, lower triangle). In addition, while the majority of the eQTLs were shared across regions, we identified 1,791 (49.6%) eQTLs with a stronger effect in one region than in any other (*lfsr* < 0.05 and > 2-fold effect size in one region compared to others). Microglia from the SVZ had the most region-specific effects with 1,045, most likely because the transcriptomic profile of this region is most distinct, followed by THA (367), STG (274), and MFG (105) (**Table S16**). We include examples of shared and region-specific eQTLs (**Figure 4C**).

To assess eQTL reproducibility and cell-type specificity, we compared MiGA eQTLs with four other external eQTL datasets, including microglia^45^, monocytes^46,47^, and bulk brain dorsolateral prefrontal cortex (DLPFC)^48^ using the q-value π1 metric^49^. We found that eQTL sharing was both cell type- and region-dependent (**Figure 4D**), with the highest sharing between MiGA and the Young et al. microglia (π1 = 0.81-0.86), but with a lower sharing in the SVZ (0.51). Sharing with monocyte eQTLs was generally slightly lower than with microglia, and sharing with bulk DLPFC eQTLs was lowest. Together, these results highlight shared genetic regulation between microglia and monocytes, which is only partly captured in whole-tissue brain data^21^.

We performed a cross-study eQTL meta-analysis (MiGA, Young^45^, MyND^47^, and Fairfax^46^) using METASOFT^50^) to assess the sharing of effects between distinct cell types. We focussed on genes associated with AD^22,51^ through METASOFT’s m-value, which is the posterior probability that the effect exists in a particular study. The comparison of m-values between microglia and monocytes showed that a large number of eQTLs in AD loci have shared effects between these two cell types, for example, *MS4A6A, RABEP1, CD33, FCER1G*, and *ABCA7*. However, there were eQTLs with specific microglial effects compared to monocytes (e.g., *BIN1, PICALM, USP6NL* and *GNGT2***; Figure 4E**). The *USP6NL* gene is an example of an eQTL with a strong effect in MiGA but not in monocytes or Young et al. (**Figure 4F**). Generally, directions of effect between monocytes and microglia were concordant (**Figure S11**), with the exception of *CASS4*. eQTLs for *CASS4* are highly significant in both MiGA and monocytes (MyND) but with opposite directions of effect (**Figure 4G**).

### Genetic regulatory effects in microglia mediate neurological disease association

We next used our meta-analyzed expression and splicing QTLs to explore whether disease-associated genetic variants may potentially act through microglia eQTL or sQTL. We downloaded publicly available GWAS summary statistics for AD^22,51–53^, PD^54^, SCZ^55^, BD^39^, and Multiple Sclerosis (MS)^56^.

We assessed colocalization between GWAS variants and QTLs using the coloc R package^57^. We compared our MiGA QTLs to the same set of published microglia, monocytes, and bulk brain tissue as before. AD and PD had the highest number of colocalizing loci in each QTL dataset, compared to the other diseases (**Figure 5A, Table S17**), with 10-30% of loci containing at least one colocalized gene, depending on the stringency of the H4 posterior probability (PP4), with lower proportions observed in BD, SCZ and MS. Within each disease the number of colocalized loci was similar between QTL datasets, although notably there were no MiGA eQTLs colocalizing in BD at PP4 ≥ 0.5.

**Figure 5.**
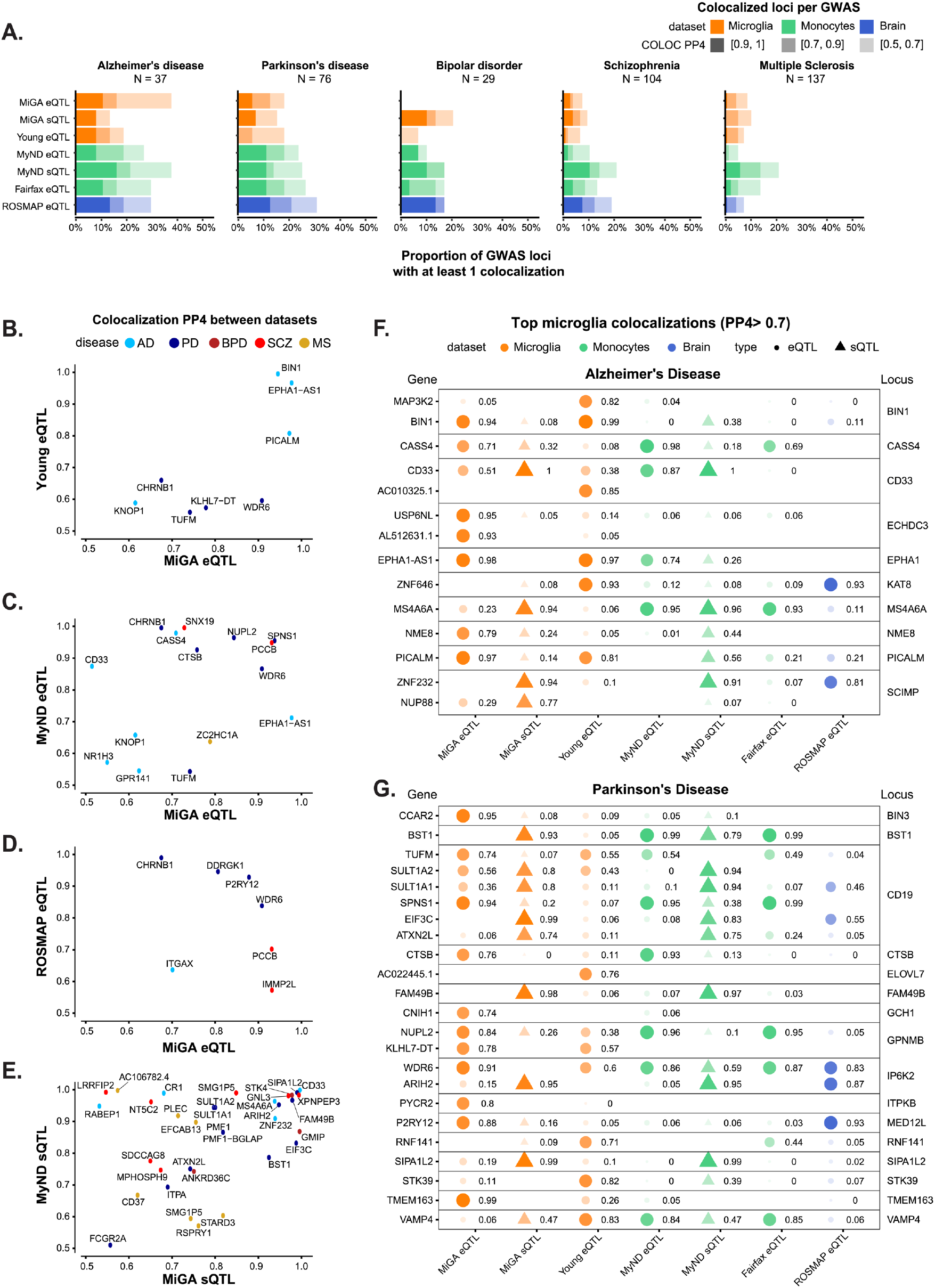
Summary of colocalization analyses. A) The proportion of GWAS loci that have at least one colocalized gene in each QTL dataset. Fill opacity used to represent numbers of loci at different stringency levels for colocalization posterior probability H4 (PP4): 0.5-0.7 (lightest), 0.7-0.9 (medium); 0.9-1 (darkest). Bars are colored by the cell or tissue type of the QTLs: microglia (orange), monocytes (green), or dorsolateral prefrontal cortex (DLPFC; blue). B-E) Pairwise comparison of the coloc PP4 for genes between QTL datasets. Points are colored by disease. B), C), D) compare the MiGA microglia eQTLs to the Young et al. microglia eQTLs, MyND monocyte eQTLs, and ROSMAP DLPFC eQTLs, respectively. E) Comparison of the MiGA splicing QTLs and the MyND monocyte splicing QTLs. F) All genes in the AD GWAS that have a PP4 > 0.7 in one of the three microglia QTL datasets. Shape opacity and size scaled to the magnitude of PP4. Circles represent colocalizations with expression QTLs and triangles represent those with splicing QTLs. G) The same for Parkinson’s Disease.

We then compared different QTL datasets to find shared evidence of colocalization at the level of individual genes within a GWAS locus (**Figure 5B-E**). The sharing between our microglia and previously published microglia^45^ was low (**Figure 5B**), with only a few known loci in AD and PD (*BIN1, PICALM, CHRNB1*), presumably due to lower power in the Young et al. data compared to our multi-tissue meta-analysis. Overall, 11% of MiGA eQTL colocalizations could be reproduced in the Young et al. data, and 15% of the Young et al. colocalizations could be found in the MiGA data, at a relaxed PP4 > 0.5, whereas sharing between the two monocyte datasets^46,47^ was 17-24% with the same parameters (**Figure S12**). Substantially lower sharing (18-21%) was observed between the MiGA eQTLs and those of our lab’s recent monocyte dataset (**Figure 5C; Figure S12**) than between the respective splicing QTLs (23-53%) **(Figure 5E; Figure S12)**. This suggests that splicing QTLs are less cell type-specific, presumably due to the association with distinct types of regulatory elements.

We present colocalizations in AD (**Figure 5F**) and PD (**Figure 5G**) in each QTL dataset. We emphasize microglia by including only genes that colocalize with one of the three microglia QTL datasets at PP4 > 0.7. In each disease there are genes that appear to be microglia-specific (*BIN1, PYCR2*), shared between microglia and monocytes (*CASS4, CTSB*), and shared between microglia and brain (*ZNF646, P2RY12*). We also observe multiple splicing QTLs, some previously reported (*CD33, FAM49B*), and some novel to this study (*MS4A6A, BST1*). We present full plots for all colocalizations in each disease in the supplementary materials (**Figures S13-22**).

### Neurological disease variants regulate gene expression in microglia and map to microglia-specific enhancers

We next examined whether the microglia eQTLs that colocalized with disease GWAS loci were due to genetic variation within microglia-specific regulatory regions. We combined the results of four different statistical and functional fine-mapping approaches to create a set of fine-mapped variants at each disease locus (see Methods, **Table S18**). We defined variants prioritized by at least one fine-mapping tool at a posterior probability ≥ 95% of being causal for the phenotype as “credible set SNPs”, and variants prioritized by at least two tools as “consensus SNPs”. In addition, because coloc does not take linkage disequilibrium (LD) structure into account, which can confound results due to non-independence between SNPs, we calculated the LD between each lead QTL variant and the set of fine-mapped SNPs at each locus using 1000 Genomes (phase 3) European reference samples (**Figure S23**). In GWAS loci with multiple colocalized genes this can be used to prioritize the most likely candidate. In the CD19 locus in PD, this approach gave additional weight to suggesting an eQTL in *SPNS1* as the likely causal gene, due to high LD between the *SPNS1* lead QTL SNP and the GWAS SNP (*r*^*2*^*=0*.*78)*, compared to *TUFM* and *SULT1A2* (**Figure S24**). With the expanded set of SNPs at each locus, we then overlapped each SNP with sets of defined promoter and enhancer regions in microglia, neurons, oligodendrocytes, and astrocytes^25^. We found that 10 out of 17 genes colocalizing in AD, 8 out of 18 in PD, 4 out of 9 in SCZ, and 3 out of 17 in MS include SNPs that overlap with microglial enhancers (**Figure 6A; Figure S25**). This approach allowed us to prioritize disease loci that likely act on disease risk by modulating gene expression specifically in microglia. Here we discuss two examples.

**Figure 6.**
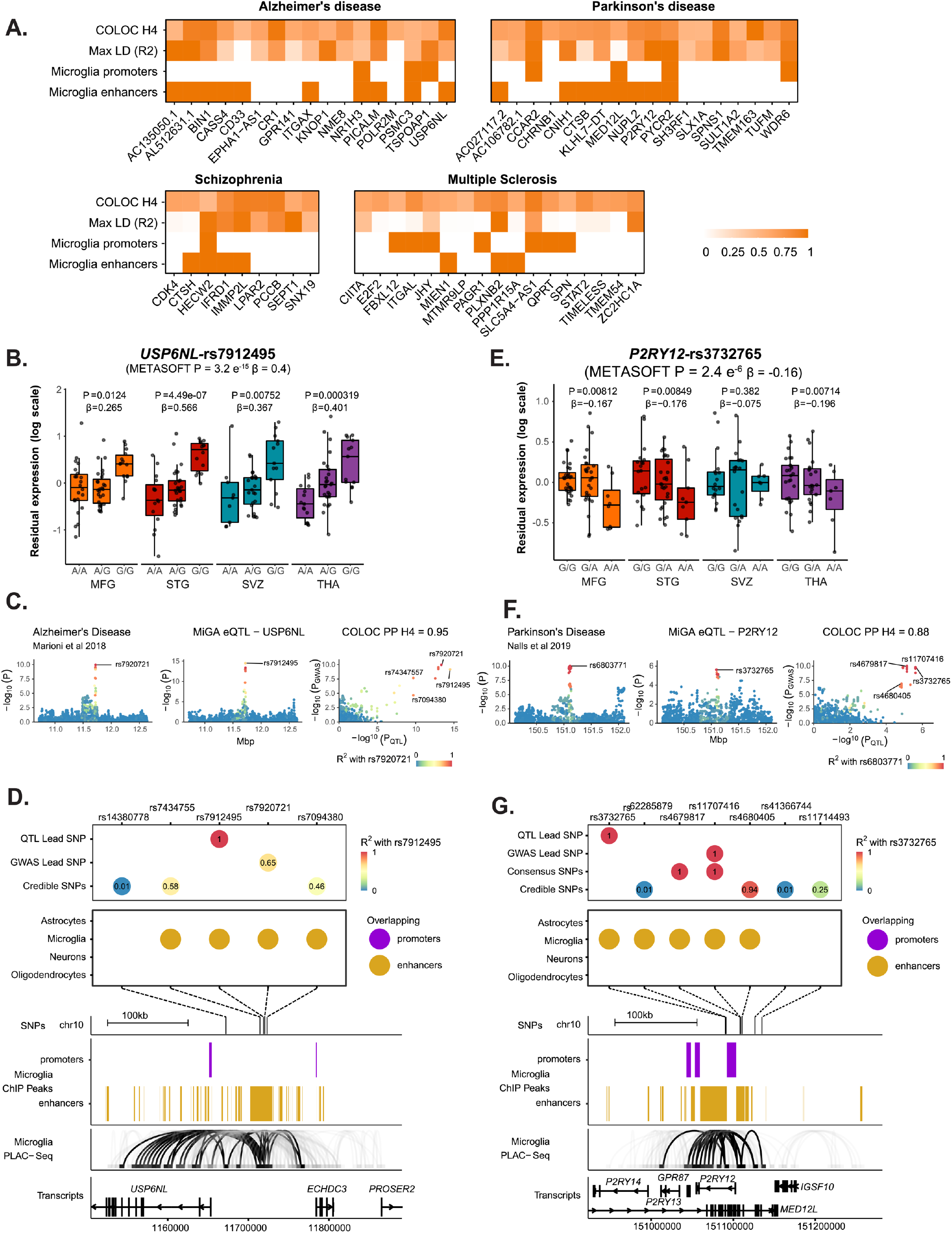
Enhancer-promoter interaction data links GWAS variants to microglia-specific regulatory regions. A) Overview of fine-mapping and epigenomic overlap analyses for all eQTL genes with PP4 > 0.5 in each disease. Max linkage disequilibrium (LD) refers to the highest LD coefficient between the lead eQTL SNP and any of the lead GWAS SNP or fine-mapped SNPs. Microglia enhancers and promoters refer to whether any of the SNPs for that eGene overlap a microglia enhancer or promoter, as defined by Nott et al. B-D) Analysis of the *USP6NL* gene. B) *USP6NL* expression is associated with the rs7912495 genotype in all four microglia regions. C) The meta-analyzed *USP6NL* eQTL colocalizes with the ECHDC3 Alzheimer’s disease risk locus (PP4 = 0.95). D) Fine-mapping of the ECHDC3 locus and combining with the lead QTL and lead GWAS SNPs. SNPs are colored by the LD with the lead QTL SNP. 4 out of 5 of the SNPs overlap a microglia-specific enhancer element as defined by ChIP-seq. Genomic plots (hg19) of the fine-mapped SNPs and the epigenomic data from microglia ChIP-seq, and PLAC-seq junctions. Junctions that overlap the fine-mapped SNPs are emphasized. E-G) Analysis of the *P2RY12* gene. E) P2RY12 expression is associated with the rs3732765 genotype. F) The *P2RY12* eQTL colocalizes with the MED12L Parkinson’s Disease locus. G) Fine-mapping of the MED12L locus discovers SNPs that in strong LD with the eQTL lead SNP that overlap microglia-specific enhancer regions. Genomic plots show that the microglia enhancer connects with the *P2RY12* promoter.

#### USP6NL

The ECHDC3 locus has been associated with AD risk in several GWAS^22,51,53^. The lead SNP rs7920721 sits in an intergenic region that separates two genes, *ECHDC3* and *USP6NL*. Both genes are interesting candidates for AD risk. eQTLs in *ECHDC3* have been associated with insulin resistance in adipose tissue^58^. *USP6NL* is a Rab GTPase activating protein^59^ involved in the retrograde transport from the Golgi apparatus^60^. Both lipid metabolism and endocytosis are pathways linked to AD through multiple risk genes^26,41,61^. Previous analyses have prioritized *ECHDC3*, as it is upregulated in AD post-mortem brains^62,63^, and an eQTL for *ECHDC3* was seen in whole blood^64^, though it did not colocalize with the GWAS SNP^22^.

*USP6NL* harbors an expression QTL observed in all four microglial regions, with the lead QTL SNP rs7912495-G increasing *USP6NL* expression (**Figure 6B**). The meta-analyzed eQTL colocalizes with the ECHDC3 locus in all 4 AD GWAS used in this study, with the highest PP4 (0.95) seen in^51^ (**Figure 6C; Figure S13**). No colocalization was observed in any other QTL dataset, though we note that *USP6NL* is expressed five-fold higher in microglia than in monocytes (MiGA median TPM = 15.77; MyND^47^ monocyte median TPM = 3.13). Fine-mapping of the ECHDC3 locus suggested three additional SNPs as well as the lead GWAS SNP (rs7920721) and lead QTL SNP (rs7912495). The GWAS lead SNP and the QTL lead SNP are in moderate LD (*r*^*2*^ *=* 0.65), as are two of the three fine-mapped SNPs (**Figure 6D)**. Of the five SNPs of interest, 4 of them overlap a microglia-specific enhancer. Using proximity ligation-assisted ChIP-seq (PLAC-seq) data^25^, we observed that the overlapping microglia enhancer region has extensive long-range connections to regions overlapping the *USP6NL* promoter and gene body. Notably, there was no colocalization of the upstream *ECHDC3* gene in any tested cell type, suggesting that *USP6NL* is the AD risk gene at this locus. The lead QTL SNP rs7912495-G increases AD risk (β = −0.0492; *P* = 6.8×10^-10^; ^51^) and we propose that it achieves this through upregulating *USP6NL* expression in microglia.

#### P2RY12

The MED12L locus was identified in the latest PD GWAS^54^. The lead SNP rs11707416 sits within a large intron of the *MED12L* gene, which overlaps with several smaller genes, one of which is *P2RY12*. A previous study prioritized *P2RY12* at this locus due to an overlap with eQTLs in blood and brain^65^.

*P2RY12* is an eQTL (lead SNP rs3732765) identified in the METASOFT meta-analysis, with the lead QTL SNP rs3732765-A decreasing *P2RY12* expression (**Figure 6E**). The eQTL colocalizes with the PD GWAS MED12L locus (PP4 = 0.88; **Figure 6F**). Colocalization was also observed with a *P2RY12* eQTL in the dorsolateral prefrontal cortex (PP4 = 0.93; **Figure 5D; Figure S15**). Fine-mapping revealed that the lead GWAS SNP rs11707416 was suggested as a causal SNP by multiple fine-mapping tools (a consensus SNP) and is in perfect LD (*r*^*2*^ *=* 1) with the lead QTL SNP rs3732765 (**Figure 6G**). In addition, there are 4 other SNPs prioritized by fine-mapping, 2 of which were in perfect or very high LD with the lead QTL SNP. Of the 7 total SNPs in the set, 5 overlapped a microglia-specific enhancer region on either side of the *P2RY12* promoter. PLAC-seq revealed long-range connections between the enhancer and the *P2RY12* promoter but not to *MED12L* (**Figure 6G**). No colocalization was observed with any *MED12L* QTL. Altogether this suggests that *P2RY12* is the causal gene at the locus. The lead QTL SNP rs3732765-A decreases PD risk (β = −0.06; *P* = 2.4×10^-10^; ^66^), and we propose that it acts through downregulating P2RY12 expression in microglia.

## Discussion

We present the largest microglia genomic resource to date. We demonstrate that transcriptional heterogeneity in human microglia varies between brain regions and across aging. We generated a catalog of eQTLs and sQTLs in microglia and identified novel disease genes and putative causal variants underlying risk for neurodegenerative and psychiatric diseases.

Regional and age-related differences in microglial density, morphology, gene, and protein expression have previously been described for both animals and humans^5,6,32,67,68^. Our current analysis across four brain regions is the largest regional- and age-related human microglia study to date and confirms transcriptional differences between cortical and subcortical microglia. Subcortical microglia show resemblances to the transcriptional response of microglia to *in vitro* culture, as well as to changes in microglia from Alzheimer’s cases. These data support the regional heterogeneity of human microglia, which may be relevant for age- and disease-related changes. Our analyses suggest some novel pathways that may be involved in regulating this regional heterogeneity, such as reelin, interferon and glucocorticoid signaling pathways. In addition, we found age-related changes in genes involved in a wide range of inflammatory responses, in line with previous results in aging in microglia^15,45,69^ and peripheral blood^43^. Of interest are a downregulation of *C2, P2RY12*, and *P2RY13*, since these molecules are emerging as key players in microglia-neuron interactions^70,71^. In addition, we found an association with genes related to two age-related disorders, AD and PD: *MS4A4A, MS4A6A*, and *BST1* are upregulated, and *P2RY12* downregulated with age. Our pathway analyses identified immune-related pathways that may be of relevance for the mechanisms of microglial aging, including STAT-3 and IL-6 signalling. Of interest is the highly significant association of LXR/RXR activation, which has emerged as a key player in regulating cholesterol homeostasis and inflammation in the brain with a potential role in neurodegenerative disorders^72–74^. Based on previous studies in humans and mice^6,9^, we expected to find region-specific patterns of age-related changes in microglia. *MS4A6A* was one of the genes that showed a region-specific effect of age^75,76^. Genetic variants for AD are related to decreased expression of *MS4A6A* in blood^75^ and monocytes^21,77^, but increased *MS4A6A* expression levels of this gene are found in AD patient blood compared to controls^75^. Our results are suggestive of a potential interaction between age and genetic risk in cortical microglia specifically.

By mapping both expression and splicing QTLs in human microglia we have created a resource that has informed our own genetic studies and will be useful for the genetics and neuroscience community. While other studies have used enhancer activity to link a GWAS lead variant to a gene^25^, or used monocyte^21,78,79^ or bulk brain eQTLs^41,48,80^, ours is the first study to link eQTLs, enhancer activity, and disease risk in microglia, and thus proposing the molecular mechanisms of action of a number of functional variants. We have identified specific disease colocalizations that may not be captured in monocytes or bulk brain tissue, like *BIN1, USP6NL*, and *PICALM* in AD, *P2RY12* in PD, *PLXNB2* in MS, and *IFRD1* in SCZ. We also found colocalizations with opposing effects, such as *CASS4*. Disease-associated eQTLs results were partly shared between MiGA and the microglia eQTL study by Young *et al*^45^. Differences between the studies in age and diagnoses of the included donors, brain regions, status (surgical versus autopsy) and sample size (93 individuals/samples versus 90 individuals/216 samples) have likely contributed to a lack of sharing of part of the hits. By mapping sQTLs we have added novel disease associations that may act through splicing, such as *MS4A6A* in AD, *SIPA1L2* and *FAM49B* in PD, *IRF3* in SCZ, *STK4 and GMIP* in BD, and *CD37* and *EFCAB13* in MS. Interpretation of these sQTLs will be improved with the generation of long-read RNA-seq in microglia to identify novel transcripts.

We have performed comprehensive fine-mapping of GWAS loci in five diseases through an ensemble of four different methods and microglia-specific epigenomic datasets to identify credible sets of putative causal variants. This approach allowed us to identify candidate functional variants in multiple disease susceptibility loci that modulate microglia-specific enhancer activity and regulate causal gene expression, which in turn likely modify disease risk by altering the function of microglia (or other myeloid cells) in the brain. In Alzheimer’s disease, we propose *USP6NL* to be the causal gene in the *ECHDC3* locus, due to both a convincing colocalization with AD GWAS and eQTL and the overlap of fine-mapped putative causal SNPs within a defined microglia enhancer which connects with the *USP6NL* promoter. *USP6NL*, a GTPase-activating protein involved in control of endocytosis, adds to a growing list of genes (*BIN1, PICALM, RABEP1, RIN3*, and *SORL1*) that implicate the dysfunction of the myeloid endolysosomal system in AD^41,78^.

In Parkinson’s disease, we propose *P2RY12* in the *MED12L* locus through a similar mechanism. *P2RY12* is a particularly interesting gene due to the increasing body of literature on its importance for the functioning of microglia^81^, as well as the proposed link between PD and purinergic signalling^82^. *P2RY12* is one of the P2Y metabotropic G-protein-coupled purinergic receptors, which is highly expressed in microglia in comparison to other brain and myeloid cell types. *P2RY12* expression is lost upon microglia activation^83^, culture^37^ and in our analyses we have shown that expression is decreased with aging. P2RY12 has been shown to play a role in microglia migration, activation, and neuronal activity^71,84^. Further validation work is required to test whether the enhancers we prioritize with fine-mapping regulate these genes specifically in microglia.

We recognize several limitations to the current study. First, although we present the largest microglia dataset to date, our sample size is still small in comparison to the largest monocyte and brain datasets. We increased power by combining the four regions in a meta-analysis, with the caveat of not adjusting for shared donors, which will have increased our false discovery rate. Another limitation is a variety of known and unknown pre- and post-mortem factors that have an impact on the microglial transcriptome, as shown by our variance partition analyses, that we could not control for in our analyses. Future work will improve the power and possibility to control for confounders by larger meta-analyses with other microglia datasets. Secondly, our ability to detect additional disease-associated eQTLs may be obscured due to the use of bulk RNA-sequencing data. Future work with large numbers of single-cell RNA-seq profiles from many individuals creates enormous opportunities for mapping eQTLs across microglial subpopulations^28^. Although such studies may be limited due to the increased sparsity of single-cell data, which may result in a lower statistical power compared to bulk RNA-seq data^85^. Lastly, many eQTLs are conditional and only revealed after specific stimuli that change the activation state of specific cell types. Thus, mapping response-eQTLs after stimulation of specific-stimuli in primary microglia may reveal novel associations that may provide further mechanistic insights into the disease-associated variants^46,86,87^.

In summary, we have performed a comprehensive assessment of the transcriptomic landscape of human microglia from multiple brain regions. We have generated an atlas of genetic effects on the human microglia transcriptome, which allowed us to identify potential causal genes and variants underlying risk for neurodegenerative and psychiatric diseases. Our findings represent mechanistic hypotheses that can now be tested with further experimental work at both the level of individual variants and the candidate genes.

## Methods

### Human brain tissue

Post-mortem brain samples were obtained from the Netherlands Brain Bank (NBB)^88^ and the Neuropathology Brain Bank and Research CoRE at Mount Sinai Hospital. The permission to collect human brain material was obtained from the Ethical Committee of the VU University Medical Center, Amsterdam, The Netherlands, and the Mount Sinai Institutional Review Board. For the Netherlands Brain bank, informed consent for autopsy, the use of brain tissue and accompanied clinical information for research purposes was obtained per donor ante-mortem. Detailed information per donor, including tissue type, age, sex, postmortem interval, pH of cerebrospinal fluid, cause of death, diagnosis, use of medication is provided in **Table S1**.

### Microglial isolation and RNA sequencing

Brain tissue was stored in Hibernate media (Gibco) at 4 °C upon processing within 24 hours after autopsy. Microglia were isolated from four regions, including medial frontal gyrus (MFG; 77 samples), superior temporal gyrus (STG; 63 samples), thalamus (THA; 60 samples), subventricular zone (SVZ; 55 samples). Microglia were isolated as described before in detail^31–33^. In brief, brain tissue was first mechanically dissociated through a metal sieve in a glucose-potassium-sodium buffer (GKN-BSA; 8.0 g/L NaCl, 0.4 g/L KCl, 1.77 g/L Na2HPO4.2H2O, 0.69 g/L NaH2PO4.H2O, 2.0 g/L D-(1)-glucose, 0.3% bovine serum albumin (BSA, Merck, Darmstadt, Germany); pH 7.4) and supplemented with collagenase Type I (3700 units/mL; Worthington, USA) and DNase I (200 µg/mL; Roche, Switzerland) or 2% of Trypsin (Invitrogen) at 37 °C for 30 min or 60 min while shaking. The suspension was put over a 100 µM cell strainer and washed with GKN-BSA buffer in the centrifuge (1800 rpm, slow brake, 4 °C, 10 min) before the pellet was resuspended in 20 mL GKN-BSA buffer. 10 mL of Percoll (Merck, Darmstadt, Germany) was added dropwise and the tissue homogenate was centrifuged at 4000 rpm (fast acceleration, slow brake at 4 °C, 30 min). The middle layer was collected and washed with GKN-BSA buffer, followed by resuspension and centrifuging in a magnetic-activated cell sorting (MACS) buffer (PBS, 1% heat-inactivated fetal cow serum (FCS), 2 mM EDTA; 1500 rpm, 10 °C, 10 min). Microglia were positively selected with CD11b-conjugated magnetic microbeads (Miltenyi Biotec, Germany) according to the manufacturer’s protocol. Microglia were stored in RLT buffer + 1% 2-Mercaptoethanol or lysed in TRIzol reagent (Invitrogen, USA). RNA was isolated using RNeasy Mini kit (Qiagen) adding the DNase I optional step or as described in detail before^31^. Library preparation was performed at Genewiz using the Ultra-low input system which uses Poly-A selection. CD11b is also present on perivascular macrophages in the CNS. However, we have previously shown by mass cytometry that the percentage of macrophages (CD206^high^) was low^32^. Moreover, we showed in a subsample (*n* = 69 samples derived from 20 donors) of our cohort that the mean percentage of P2Y12^+^CD64^+^ cells is 96%, ranging from 87.2 to 99.9% (see **Figure S2**). We performed this study using a validated protocol for post-mortem microglia isolation^31–33,89^.

SMART-Seq v4 Ultra Low Input RNA Kit was used for library construction using 100 ng of RNA. The libraries were sequenced as 150 bp on fragments with an average read depth of 29 million (ranging from 14-82M) read pairs on the Illumina HiSeq 2500. RNA-seq data was processed using the RAPiD pipeline^90^. RAPiD aligns samples to the hg38 genome build using STAR^91^ using the GENCODE v30 transcriptome reference and calculates quality control metrics using Picard^92^. RNA-seq quality control was performed applying three filters to remove samples: 1) samples with less than 10M reads aligned from STAR; 2) samples with more than 20% of the reads aligned to ribosomal regions; 3) samples with less than 10% of the reads mapping to coding regions; 4) samples from brain regions with fewer than 20 donors. Estimated transcript abundance was obtained using RSEM^93^ and transcripts were summed to the gene level with tximport^94^. Genes with more than 1 read count per million (CPM) in 30% of the samples were kept for downstream analysis. Gene level read counts were normalized as transcripts per million mapped reads (TPM) to adjust for sequencing library size differences.

### DNA isolation and genotyping

Genomic DNA was extracted from medial frontal gyrus, superior temporal gyrus, thalamus, or cerebellum using the Qiagen DNeasy Blood & Tissue Kit and followed the manufacturer’s instructions. In short, a small piece of brain tissue (0.020-0.040 gram) was cut and placed in a 96-deepwell plate (Thermo Fisher) on dry ice. A mastermix containing binding enhancer, PBS and proteinase K (Qiagen) was added to each well. The solution was incubated overnight at 65 degrees Celsius. The KingFisher(tm) (KF) Duo system was applied with a 12 pin magnet head which enabled processing of 12 samples per run using microtiter 96 deepwell plates (Thermo Fisher). Prior to the extraction process, the deepwell plates were filled with the following reagents: wash buffer (Qiagen), 80% Ethanol, elution buffer (Qiagen) and tip combs (Qiagen). For extraction, 440 μl of DNA binding buffer beads (Qiagen) were added to the sample and mixed by pipetting. After that, automated extraction was executed and completed within 30 minutes using the protocol provided by the manufacturer. The extracted DNA was eluted in 200 μl elution buffer and subsequently transferred to 1.5 ml tubes for storage. DNA quality and concentration was assessed using a Nanodrop. Samples were genotyped using the Illumina Infinium Global Screening Array (GSA), which contains a genome-wide backbone of 642,824 common variants plus custom disease SNP content (∼ 60,000 SNPs).

### External Datasets

We downloaded genome-wide association study (GWAS) and genome-wide association study by proxy (GWAX) summary statistics for the following diseases: Alzheimer’s disease (AD)^22,51–53^, Parkinson’s disease (PD)^66^, Schizophrenia (SCZ)^55^, Bipolar Disorder (BD)^39^, and Multiple Sclerosis (MS)^56^. For each GWAS we downloaded the full summary statistics and a list of genome-wide significant loci, as defined separately by each study. Missing fields in the nominal statistics were dealt with as follows: standard error was calculated from the effect size and *P*-value; minor allele frequency was taken from the European samples from 1000 Genomes^95^; SNP coordinates or RS ID were matched using Ensembl (release 99). We took the lists of top genome-wide associated loci from the supplementary materials from each study. For the PD GWAS we removed any loci that did not pass the final quality control filtering according to the “Failed final filtering and QC” column. To avoid double-counting in colocalization, if multiple GWAS loci overlapped (within 1 megabase), we retained the locus with the lowest *P*-value. Due to the complex LD structure within both regions^96,97^, loci overlapping the human MHC/HLA region (hg19 chr6:28,477,797-33,448,354) or the MAPT H1/H2 haplotype region (hg19 chr17:43,628,944-44,571,603) were removed. When conditionally independent loci were listed, only the primary association was kept due to lack of conditional summary statistics. Loci from the four AD GWAS were given consensus names using the most recent GWAS as a guide. This resulted in the following locus numbers for each disease: 37 for AD, 71 for PD, 104 for SCZ, 29 for BD, and 137 for MS.

Expression quantitative trait loci (eQTL) full summary statistics were downloaded for microglia^27^, monocytes^46,47^ and dorsolateral prefrontal cortex^48^. All summary statistics were coordinate sorted and indexed with tabix^98^. Epigenomic data from purified human microglia, neurons, astrocytes, oligodendrocytes^25^ were downloaded through the echolocatoR package^99^.

To replicate and validate our findings we downloaded processed lists of differentially expressed genes from previous microglia studies^5,15,27^.

### Sources of transcriptomic variation

To understand major sources of variation in the gene expression data at the sample level, we used PCA and linear regression to measure the effect of the following experimental confounders on gene expression variance: sex, age, post mortem interval (PMI), pH, and technical covariates estimated by Picard **(Figure S4**). We then applied variancePartition (v1.17.7), which uses a linear mixed model to attribute a percentage of variation in expression based on selected covariates on each gene^34^. As highly correlated covariates cannot be included in the model, we selected covariates that were not very strongly correlated to run the variancePartition analysis (**Figure S5B**). Gene counts were normalized using trimmed means of M-values (TMM) values calculated from edgeR^100^ and *voom* transformed^101^, which is a method that estimates the mean-variance relationship of the log-counts as input to variancePartition. The technical covariates included in the analysis were % mRNA bases (Picard), mean insert size (Picard), % ribosomal bases (Picard), % read alignment (Picard), and sequencing lane. The biological covariates were donor ID, donor age, sex, brain region, cause of death, sample pH, main diagnosis, post-mortem interval (PMI) in minutes, and the first 4 genotyping ancestry MDS values (C1-C4).

### Differential Expression Analysis

Differential expression analysis was performed between the brain regions using the R package Differential expression for repeated measures (DREAM) from VariancePartition^35^. DREAM uses a linear model to increase power and decrease false positives for RNA-seq datasets with repeated measurements. For the analysis, inputs included the count matrix and the covariate file. These data were normalized using the function voomWithDreamWeights that also performs voom transformation. Since one donor can have different brain regions, we modeled the individual as a random effect and added selected covariates to adjust for possible technical and biological confounders. The final model accounted for sex, donor ID, age, region, cause of death, the first 4 ancestry MDS values (C1-4), % mRNA bases, median insert size, and % ribosomal bases. *P*-values were then adjusted for multiple testing correction using the Benjamini-Hochberg False Discovery Rate (FDR) correction.

Differential age-by-region gene expression was measured by fitting linear mixed models using DREAM (considering repeated donor measures). Analogously as in the standard region differential analysis, expression data were first normalized and transformed using voomWithDreamWeights. Four models were fitted, setting each brain region as a reference. The model included sex, donor ID, cause of death, the first 4 ancestry MDS values (C1-4), % mRNA bases, median insert size, and % ribosomal bases, plus an interaction term with age and region. Next, we extracted the interaction coefficients for all pairwise region comparisons and selected genes with FDR < 0.05.

### Pathway and Gene Set Enrichment analysis

#### Pathway analysis

we performed canonical pathway analyses in the Ingenuity Pathway Analysis (IPA) software independently using the following input gene sets: upregulated DEGs aging (n = 338 genes), downregulated DEGs aging (n = 1,355 genes), and clusters of gene sets for specific brain regions; cluster 1 (n = 333 genes upregulated in MFG and STG), cluster 2 (n = 108 genes upregulated in SVZ and THA), cluster 3 (n = 350 genes downregulated in SVZ), and cluster 4 (n = 296 genes upregulated in SVZ) at FDR < 0.05. We show the top 10 enriched significant pathways in **Table S9** and **S11**. Three to five out of the 10 significant enriched pathways, specifically related to microglia function, with at least four genes that overlap are shown in **Figures 2F and 3B**.

#### Gene set enrichment analysis

to test specific pathways we used curated gene sets and tested statistical enrichment using Fisher exact test at FDR < 0.05 for the following curated gene lists: (1) Human Alzheimer disease (HAM) curated lists: 53 upregulated and 22 downregulated genes from^38^ (2) Cultured microglia curated lists: raw counts were extracted from^37^. The Bioconductor package DESeq2^102^ was employed to determine differential gene expression between ex vivo and microglia samples cultured for 7 days; 3,674 upregulated and 4,121 downregulated genes were detected and used in further analyses. (3) IFN-y stimulated microglia curated gene list: 74 upregulated and 6 downregulated genes were detected following the methods as described below in IFNy and LPS stimulated microglia (4) LPS stimulated microglia curated genes list: 472 upregulated and 316 downregulated genes were detected following the methods as described below in IFN-y and LPS stimulated microglia. (5) Aging in human peripheral blood curated gene list: 600 upregulated and 897 downregulated genes from^43^. (6) Microglia-specific curated list: 249 genes from^103^. Additionally, we included specific disease-related lists based on the latest TWAS results: (10) Alzheimer’s disease curated gene list: 36 genes from^41^ (11) Parkinson’s disease curated gene list: 77 genes from^42^ (12) Schizophrenia curated gene list: 43 genes from^40^ (13) Bipolar disorder curated gene list: 16 genes from TWAS results from latest BD GWAS^39^. For curated gene lists from mouse datasets, we used g:Convert, a gene identifier conversion tool to convert mouse to human ensembl IDs^104^.

### IFN-y and LPS stimulated microglia

Microglia were isolated from the MFG of 6 unaffected controls, cultured and stimulated as described before^31,32^. In summary, after overnight incubation, microglial medium was supplemented with 100 ng/mL LPS from Escherichia coli 0111:B4 (Merck, Germany) or with 50 ng/ml interferon gamma INF-γ for 6 hours. Microglial RNA was isolated using the TRIzol method and cDNA libraries were generated using the SMART-Seq v4 Ultra Low Input RNA Kit for Sequencing Components (Takara) according to the manufacturer’s protocol.The libraries were sequenced as 150 bp paired-end reads with an average read depth of 42 million (range 16-98M) read pairs on an Illumina HiSeq 2500. FASTQ files were processed through RAPiD as described above. Two outliers (one unstimulated sample and one LPS-exposed sample) were detected with the hclust function in R and excluded from further analysis. Differential expression was tested using DESeq2^102^, testing all genes with greater than 1 read count per million in at least 30% of samples. The effect of condition was calculated separately for LPS and IFNy, controlling for differences in donor in both cases. Since donor highly correlated with all other covariates measured, no other variables were included in the design. Genes with an adjusted P-value of < 0.05 were considered significant.

### Genotype Quality Control and Imputation

Samples were genotyped using the Illumina Infinium Global Screening Array (GSA) plus a custom disease SNP content (∼ 60,000 SNPs) for a total of 760,329 common variants. To select high-quality data, we applied an initial genotyping quality control, keeping SNPs with call rate > 95%, minor allele frequency (MAF) > 5%, Hardy-Weinberg equilibrium (HWE) *P*-value > 1 x 10^-6^, and sample call rate > 95%.

Duplicated and up to third-degree related samples were removed based on pairwise kinship coefficients estimated using KING^105^ (**Figure S9A**). DNA samples were matched to the RNA-seq data to confirm the same donor origin using the MBV tool from QTLtools^106^ (**Figure 9B**) and sex mismatching samples were removed by comparing DNA inferred sex from PLINK to RNA gene expression of the *UTY* and *XIST* genes (**Figure S9C**). This resulted in 593,748 genotyped variants passing all QC steps in 98 donors, of which 90 donors were of European ancestry. Genetic ancestry of samples was confirmed by principal components analysis using the PLINK program^107^ and MDS (multidimensional scaling) values of study subjects were compared to those of 1000 Genome Project samples (Phase 3) (**Figure S9D**).

Genotype imputation was performed for those 90 donors through the Michigan Imputation Server v1.4.1 (Minimac 4)^108^ using the 1000 Genomes (Phase 3) v5 (GRCh37) European panel and Eagle v2.4 phasing^109^ in quality control and imputation mode with rsq filter set to 0.3. Following imputation, variants were lifted over to the GRCh38 reference to match the RNAseq data using Picard liftoverVCF and the “b37ToHg38.over.chain.gz” liftover chain file. Finally, we applied another round of variant quality controls, removing indels and multi-allelic SNPs, and keeping only variants with MAF > 5% and Hardy-Weinberg *P*-value >1×10^-6^. After imputation, liftover, and QC, a total of 5,803,004 variants were included in downstream analyses. These variants were additionally annotated using dbSNP (All_20180418.vcf.gz) and snpEff v4.3i^110^.

### Quantitative Trait Loci mapping

To perform expression QTL (eQTL) mapping, we followed the latest pipeline created by the GTEX consortium^111^. We completed a separate normalization and filtering method to previous analyses. Gene expression matrices were created from the RSEM output using tximport^94^. Matrices were then converted to GCT format, TMM normalized, filtered for lowly expressed genes, removing any gene with less than 0.1 TPM in 20% of samples and at least 6 counts in 20% of samples. Each gene was then inverse-normal transformed across samples. After filtering, we tested a total of 18,430 genes. Then, PEER^112^ factors were calculated to estimate hidden confounders within our expression data. We created a combined covariate matrix that included the PEER factors and the first 4 genotyping ancestry MDS values as input to the analysis. We tested numbers of PEER factors from 0 to 20 and found that between 5 and 10 factors produced the largest number of eGenes in each region (**Figure S10**).

To test for *cis*-eQTLs, linear regression was performed using the tensorQTL^113^ cis_nominal mode for each SNP-gene pair using a 1 megabase window within the transcription start site (TSS) of a gene. To test for association between gene expression and the top variant in *cis* we used tensorQTL cis permutation pass per gene with 1000 permutations. To identify eGenes, we performed q-value correction of the permutation *P-*values for the top association per gene^49^ at a threshold of 0.05.

We performed splicing quantitative trait loci (sQTL) analysis using the splice junction read counts generated by regtools^114^. Junctions were clustered using Leafcutter^115^, specifying for each junction in a cluster a maximum length of 100kb. Following the GTEx pipeline, introns without read counts in at least 50% of samples or with fewer than 10 read counts in at least 10% of samples were removed. Introns with insufficient variability across samples were removed. Filtered counts were then normalized using prepare_phenotype_table.py from Leafcutter, merged, and converted to BED format, using the coordinates from the middle of the intron cluster. We created a combined covariate matrix that included the PEER factors and the first 4 genotyping ancestry MDS values as input to the analysis. We mapped sQTLs with between 0 and 20 PEER factors as covariates in our QTL model and determined 5 to be optimal in MFG, STG and THA. 0 PEER factors were used for SVZ (**Figure S11**).

To test for cis sQTLs, linear regression was performed using the tensorQTL nominal pass for each SNP-junction pair using a 100kb window from the center of each intron cluster. To test for association between intronic ratio and the top variant in *cis* we used tensorQTL permutation pass, grouping junctions by their cluster using --grp option. To identify significant clusters, we performed q-value correction using a threshold of 0.05.

We estimated pairwise replication (π1) of cis-eQTLs with the external eQTL datasets using the q-value R package^49^. Briefly, this involves taking the SNP-gene pairs that are significant in our microglia data at q-value < 0.05 and extracting the unadjusted *P*-values for the matched SNP-gene pairs in the external dataset.

### Meta-analysis of microglia QTLs

#### METASOFT

Meta-analysis of the four microglia brain regions (MFG, STG, THA and SVZ), along with monocytes (MyND and Fairfax) and dorsolateral prefrontal cortex (ROSMAP) was performed using METASOFT^50^. Effect sizes and standard errors of each SNP-Gene pair were used as input. We carried out a random effects meta-analysis using their RE2 model, optimized to detect associations under heterogeneity.

#### mashr: Multivariate Adaptive Shrinkage in R

To estimate and compare the genetic effects in gene expression and splicing proportions across different brain regions, we performed a Multivariate Adaptive Shrinkage (MASH) through the R package mashR^44^. MASH employs an empirical Bayes method to estimate patterns of similarity among conditions and improve the accuracy of effect estimates.

Following the pipeline applied by GTEx Consortium (see URLs) we used as input, the nominal associations (P-values, betas, and standard errors) from eQTL and sQTL (gene-SNP pair for eQTL or junction-SNP pair for sQTL) for each region. Then, the analysis was performed as follows:

1. Select the strongest associations after computing a sparse factorization matrix of the z-scores using the Sparse Factor Analysis (SFA) software with K=5 factors.
2. Compute data-driven covariance matrices priors by applying the Extreme Deconvolution method and compute the canonical covariance matrices, including the identity matrix, and matrices representing condition-specific effects.
3. Next, using the entire dataset, compute the maximum-likelihood estimates of the weights for each combination and learn how each pattern-effect size combination occurs in the data.

Finally, we computed the posterior statistics using the fitted MASH model from the previous step. This step creates the tables with posterior means and local false sign rate (*lfsr*), a measure analogous to FDR, that accounts for effect size and standard error rather than only *P*-values^116^. This approach improves effect size estimates and allows for more quantitative assessments of effect-size heterogeneity compared to simple region-specific assessments^44^.

### Colocalization and fine-mapping

For the MiGA eQTLs and sQTLs we used the METASOFT random effects meta-analysis across the four regions for colocalization analyses. We used the coloc package^57^ to test whether SNPs from different disease GWAS colocalized with expression and splicing QTLs from microglia, monocytes and brain (dorsolateral prefrontal cortex). For each genome-wide significant locus in a GWAS we extracted the nominal summary statistics of association for all SNPs within 1 megabase either upstream/downstream of the top lead SNP (2Mb-wide region total). In each QTL dataset we then extracted all nominal associations for all SNP-gene pairs within that range and tested for colocalization between the GWAS locus and each gene. We used thresholds of posterior probability H4 (PP4) ≥ 0.5 for suggestive, ≥ 0.7 for moderate and ≥ 0.9 for strong colocalization, respectively. We restricted our colocalizations to GWAS SNP - eQTL SNP pairs where the distance between their respective top SNPs was ≤ 500kb or the two lead SNPs were in moderate linkage disequilibrium (r^2^ > 0.1), taken from the 1000 Genomes (Phase 3) European populations using the LDLinkR package^117^).

For splicing QTLs we followed the same approach but collapsed junctions to return only the highest PP4 value for each gene in each locus. For presenting results across diseases, we merged overlapping loci from the four Alzheimer’s disease studies together, presenting results with the highest PP4 value for each gene. Due to the smaller window of association (100kb from the center of the intron excision cluster) we restricted reported colocalizations to cases where the GWAS SNP and the top sQTL SNP were either within 100kb of each other or in moderate linkage disequilibrium (r^2^ > 0.1).

We used echolocatoR^99^ to perform statistical and functional fine-mapping of each GWAS locus with a suggestive colocalization PP4 > 0.5. echolocatoR combines the output of multiple fine-mapping tools to identify high-confidence putative causal SNPs. The full 2Mb window was fine-mapped in each locus to better take into account more widespread LD architectures. SNPs with minor allele frequency (MAF) < 5% were removed as we were primarily focused in identifying common risk factors. We fine-mapped using ABF^118^, FINEMAP^119^, SUSIE^120^, and POLYFUN + SUSIE^121^, with 1000 Genomes (Phase 3) European samples as our LD reference. For tools that permitted, we set the maximum number of causal SNPs per locus to 5. Each tool produces a 95% credible set of SNPs, which can be understood as SNPs with a posterior probability > 95% of being causal for the given phenotype. echolocatoR then defines SNPs that are present in multiple tools’ credible sets as “consensus SNPs” with a higher level of confidence in their causality. Importantly, we define the lead GWAS SNP as the lead variant listed in the original summary statistics, which may or may be not prioritized by fine-mapping as a credible or consensus SNP.

### Cell-type specific promoter-enhancer data

We downloaded processed cell-type specific promoter and enhancer data^25^ using the echolocatoR package^99^. Briefly, fluorescent activated nuclear sorting (FANS) was performed on post-mortem human brains to isolate PU.1+ microglia, NEUN+ neurons, OLIG2+ oligodendrocytes, and NEUN-LHX2+ astrocyte nuclei. Chromatin immunoprecipitation sequencing (ChIP-seq) was performed for the histone modifications H3K27ac and H3K4me3, which identify activated chromatin regions and promoters, respectively. Regions were defined by Nott and colleagues as active promoters when a H3K4me3 peak overlapped H3K27ac within 2000bp of the nearest transcription start site, whereas active enhancers were defined as H3K27ac peaks that did not overlap any H3K4me3 peaks. In addition, they performed proximity ligation-assisted ChIP-seq (PLAC-seq^122^), which identifies long-range connections between H3K4me3-positive promoter regions and other genomic regions. We downloaded coordinates for ChIP-seq and PLAC-seq in each cell type and overlapped our SNP sets in each colocalized locus using GenomicRanges in R^123^.

### Visualization

All plots were created using ggplot2^124^ in R (version 3.6.0), with ggrepel^125^, ggfortify^126^, patchwork^127^, and ggbio^128^ for additional layers of visualization.

## Acknowledgments

We thank members of the Raj and de Witte labs for their feedback on the manuscript. This work was supported by grants from the US National Institutes of Health (NIH NIA R21-AG063130, NIA R01-AG054005, and NIA R56-AG055824). This work was supported in part through the computational and data resources and staff expertise provided by Scientific Computing at the Icahn School of Medicine at Mount Sinai. Research reported in this paper was supported by the Office of Research Infrastructure of the National Institutes of Health under award number S10OD026880. The authors thank Michael Chao for his assistance with genotyping QC. The authors thank the teams of the Netherlands Brain Bank and the Mount Sinai Neuropathology Brain Bank and Research CoRE for their services. We thank the study participants for their generous gifts of brain donation. The microglia were isolated through the efforts of a large team and we would like to thank Manja Litjens, Roland D. van Dijk, Alba Fernández-Andreu, Paul R. Ormel, Hans C. van Mierlo, Y. He, Stephanie Gumbs, Miriam E van Strien, Saskia Burm, Vanessa Donega, and Elly M. Hol for all their contributions to this effort. Gijsje Snijders was supported through ZonMw and the foundation “De Drie Lichten” in the Netherlands.

## Author Contributions

LDW and TR conceived and supervised the study. GJLS, MAMS, and ABvB isolated microglia at University Medical Center Utrecht, the Netherlands. GJLS, EN, AA, MP, FAG, and RK isolated microglia at Mount Sinai School of Medicine, New York City. EN, AA, and MP performed genotyping and RNA-seq. WvZ performed RNA-seq on stimulated microglia samples, with input from GJLS and LDW. KPL performed data pre-processing and quality control. KPL led analyses of the region, aging, QTL analyses and meta-analysis, with input from JH and GJLS. GJLS led data interpretation, functional overlaps and replication work. JH led genetic, fine-mapping and epigenomic analyses. BMS assisted with fine-mapping analyses. RAV assisted with QTL mapping and performed genotyping QC. RsK provided funding and was involved in establishing the Netherlands Brain Bank for Psychiatry (NBB-PSY), providing tissue for this project. RM performed validation work. JP and CB provided data for validation. The manuscript was written by JH, GJLS, KPL, LDW, and TR, with input from all co-authors. All authors read and approved the manuscript.

## Data and materials availability

Raw RNA-seq data generated in this work are available through Gene Expression Omnibus (GEO) (in progress). Imputed genotype data are available from the Database of Genotypes and Phenotypes (dbGAP) (in progress). All differential expression, gene lists, and fine-mapping results are present as supplementary tables. The GWAS fine-mapping results are available from the echoLocatoR Shiny application (see URLs). Full nominal and permuted eQTL and sQTL summary statistics per brain region are available from Zenodo at https://doi.org/10.5281/zenodo.4118605 (eQTL) and https://doi.org/10.5281/zenodo.4118403 (sQTL). Results for eQTL and sQTL meta-analysis (mashR and METASOFT) and colocalization (COLOC) are available from Zenodo at https://doi.org/10.5281/zenodo.4118676.

## URLs

GTEx QTL mapping pipeline: https://github.com/broadinstitute/gtex-pipeline

MASHR example analysis of GTEX data: https://stephenslab.github.io/gtexresults/gtex.html

echolocatoR Shiny Application: https://rajlab.shinyapps.io/Fine_Mapping_Shiny

