## Supplementary Figures 1-25 for "Atlas of genetic effects in human microglia transcriptome across brain regions, aging and disease pathologies"

### List of figures

Supplementary Figure 1: Flowchart of quality control. 4

Supplementary Figure 2: Purity of microglia samples. 5

Supplementary Figure 3: Overview of the data. 6

Supplementary Figure 4: Sources of variation in the gene expression data. 7

Supplementary Figure 5: Main sources of expression variation and correlation of covariates. 8

Supplementary Figure 6: Principal component analysis (PCAs) and data adjustment. 10

Supplementary Figure 7: Replication of MiGA age-related genes with peripheral human blood. 11

Supplementary Figure 8: Differential expression by age interaction with brain regions. 12

Supplementary Figure 9: Genotyping QC. 13

Supplementary Figure 10: Probabilistic Estimation of Expression Residuals (PEER) correction for non-genetic factors in eQTL and sQTL analyses. 14

Supplementary Figure 11: eQTL effect sizes of the AD-associated genes. 15

Supplementary Figure 12: Pairwise sharing of colocalized genes. 16

Supplementary Figure 13: Full colocalization results in Alzheimer’s Disease. 18

Supplementary Figure 14: Colocalization results for each regional microglia dataset in Alzheimer’s Disease. 19

Supplementary Figure 15: Full colocalization results in Parkinson’s Disease. 21

Supplementary Figure 16: Colocalization results for each regional microglia dataset in Parkinson’s Disease. 22

Supplementary Figure 17: Full colocalization results in schizophrenia. 24

Supplementary Figure 18: Colocalization results for each regional microglia dataset in schizophrenia. 25

Supplementary Figure 19: Full colocalization results in bipolar disorder. 26

Supplementary Figure 20: Colocalization results for each regional microglia dataset in bipolar disorder. 27

Supplementary Figure 21: Full colocalization results in multiple sclerosis. 29

Supplementary Figure 22: Colocalization results for each regional microglia dataset in multiple sclerosis. 30

Supplementary Figure 23: Fine-mapping of loci colocalizing with MiGA eQTLs. 31

Supplementary Figure 24: Fine-mapping and LD calculation aids gene prioritization at the CD19 Parkinson’s Disease locus. 32

Supplementary Figure 25: Overlap of colocalized microglia eQTLs with epigenomic features. 33

### List of tables

Supplementary Tables 1-18 are available as a separate file (in .xls format). Full nominal and permuted eQTL and sQTL summary statistics per brain region are available from Zenodo at https://doi.org/10.5281/zenodo.4118605 (eQTL) and https://doi.org/10.5281/zenodo.4118403 (sQTL). Results for eQTL and sQTL meta-analysis and colocalization with diseases are available from Zenodo at https://doi.org/10.5281/zenodo.4118676.

Supplementary Table 1: Metadata for all microglia samples (n = 255).

Supplementary Table 2: Full summary statistics from microglia differential gene expression between male and female.

Supplementary Table 3: Full summary statistics from microglia differential gene expression between MFG and SVZ.

Supplementary Table 4: Full summary statistics from microglia differential gene expression between SVZ and STG.

Supplementary Table 5: Full summary statistics from microglia differential gene expression between THA and MFG.

Supplementary Table 6: Full summary statistics from microglia differential gene expression between THA and STG.

Supplementary Table 7: Full summary statistics from microglia differential gene expression between THA and SVZ.

Supplementary Table 8: Full summary statistics from microglia differential gene expression between MFG and STG.

Supplementary Table 9: Microglia K-means clusters pathway analysis results and replication with external dataset (Van der Poel et al. 2019).

Supplementary Table 10: Full summary statistics from age-related analysis (Dream software).

Supplementary Table 11: Microglia pathway analysis results for the significant age-related genes (up and down regulated at FDR < 0.05) and replication with external datasets.

Supplementary Table 12: Full summary statistics from the differential gene expression analysis using an interaction-term between age and region.

Supplementary Table 13: Number of eGenes, sGenes and sClusters per region.

Supplementary Table 14: Meta-analysis mashR significant results (lfsr < 0.05) for eQTLs.

Supplementary Table 15: Meta-analysis mashR significant results (lfsr < 0.05) for sQTLs.

Supplementary Table 16: Region-specific eQTLs according to mashR (lfsr < 0.05 and effect size at least 2-fold larger in one tissue than in any other)

Supplementary Table 17: COLOC results at PP4 > 0.5.

Supplementary Table 18: Fine-mapping results for the 79 loci that are colocalized with MiGA eQTLs.

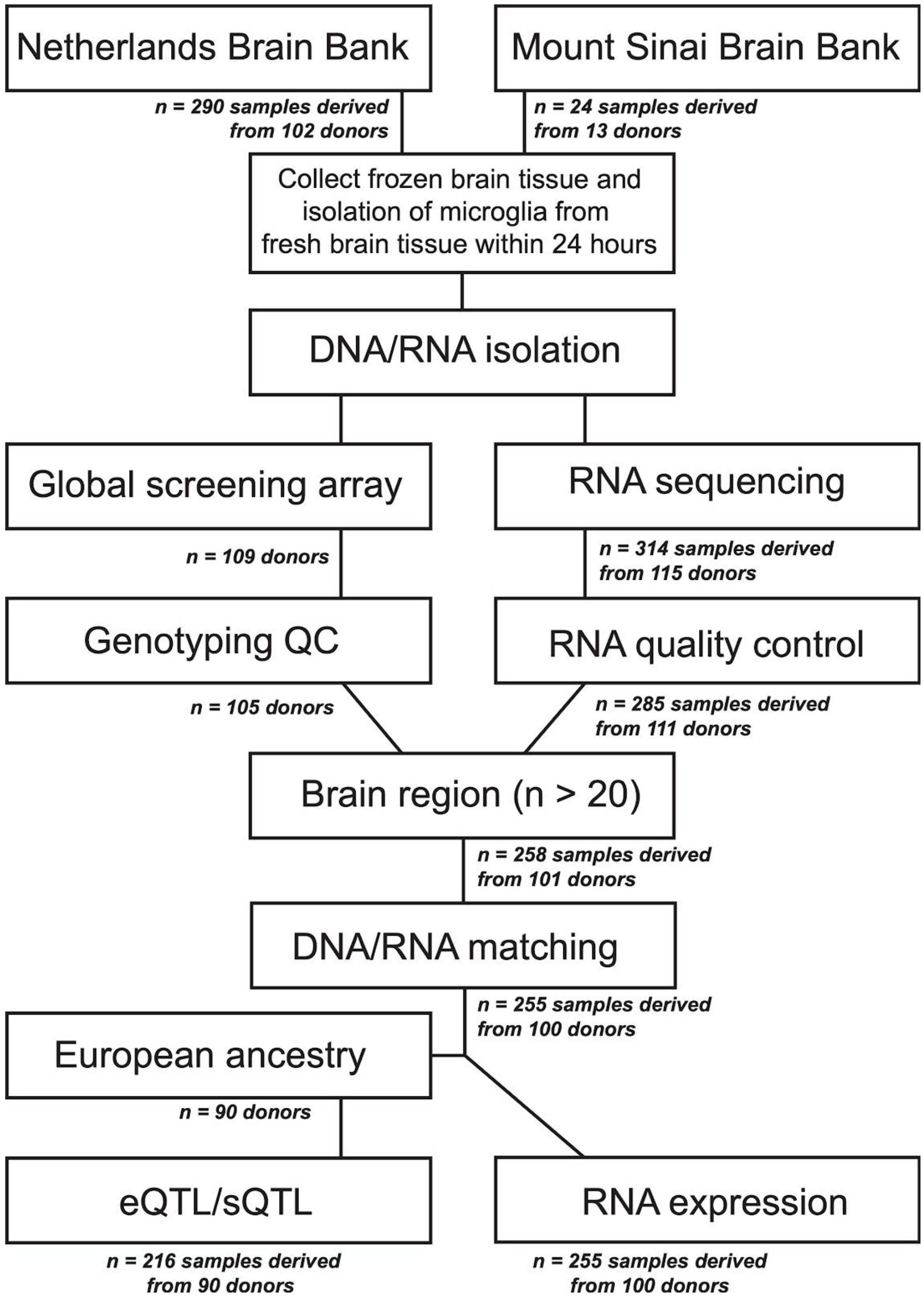


**Supplementary Figure 1: Flowchart of quality control.**

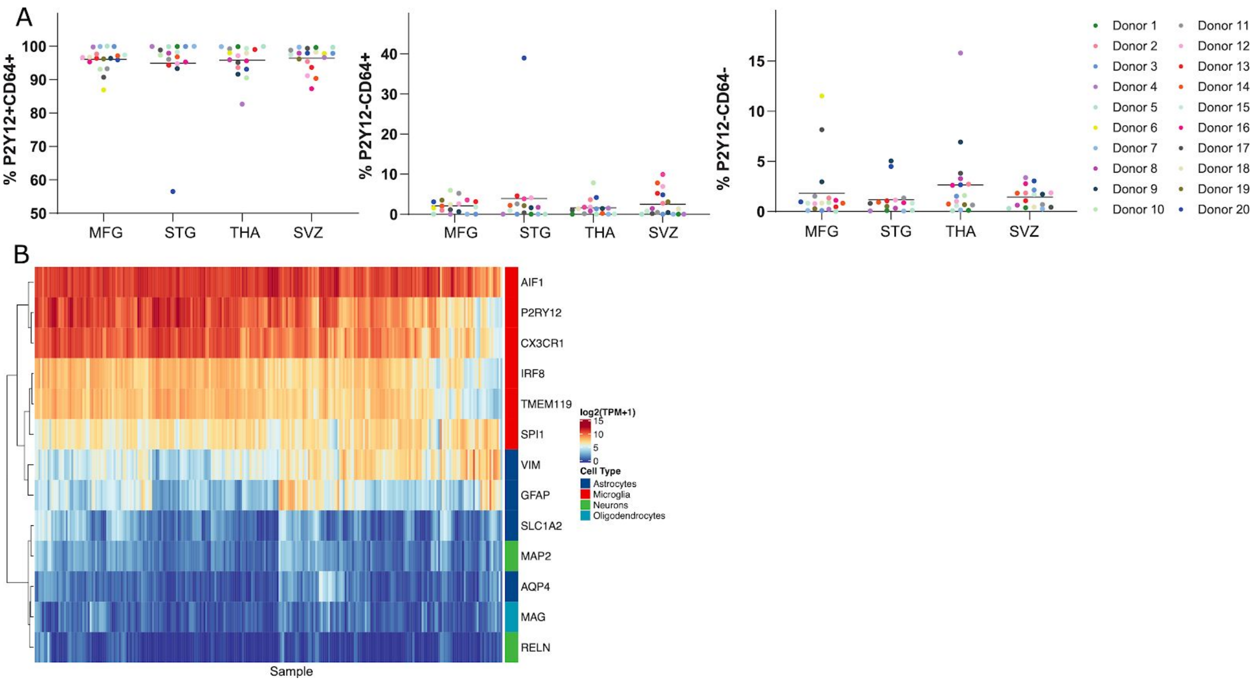


**Supplementary Figure 2: Purity of microglia samples.**

A) CyTOF analysis of CD11b-isolated cells on a subsample of *n* = 20 donors. Cells are gated for P2Y12+CD64+ (microglia); P2Y12-/CD64+ (macrophages) and P2Y12-/CD64- (non-myeloid cells) B) Expression levels (TPM+1 $log_2$ scale) of cell markers for 255 samples. Blue colors indicate low expression and red colors indicate high gene expression.

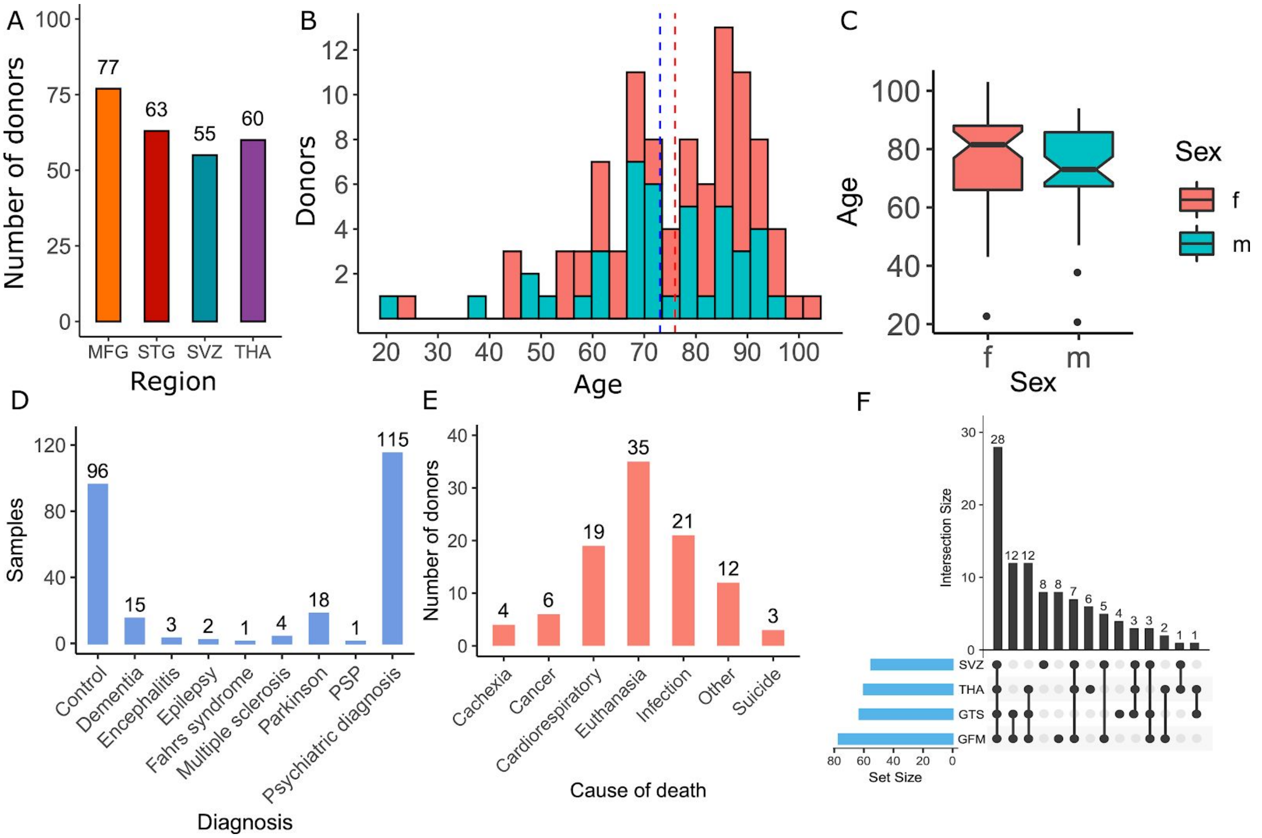


**Supplementary Figure 3: Overview of the data.**

A) Number of donors by brain region. B-C) Age range of the 100 donors in this study. Blue indicates the distribution of male donors and red indicates the distribution of female donors. Dashed lines for mean age. D) Frequency of diagnosis. E) Frequency of cause of death. F) The number of brain regions in this study. Each donor donated one up to four brain regions.

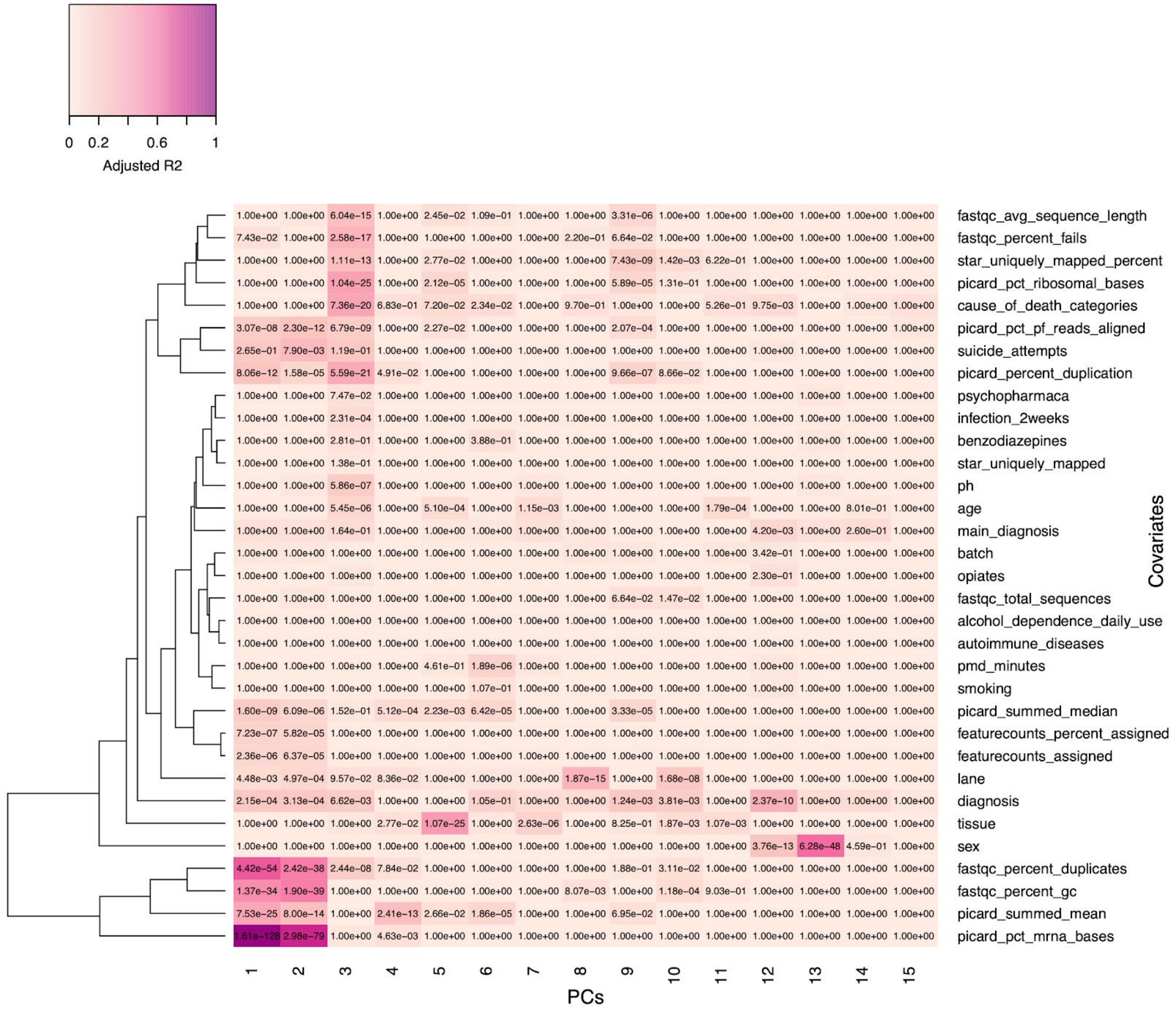


**Supplementary Figure 4: Sources of variation in the gene expression data.**

Linear regression between the first 15 Principal Components (PCs) and the covariates. Colors correspond to the adjusted R-squared and the *P*-values are Bonferroni adjusted. “Tissue” refers to which brain region the microglia were isolated from.

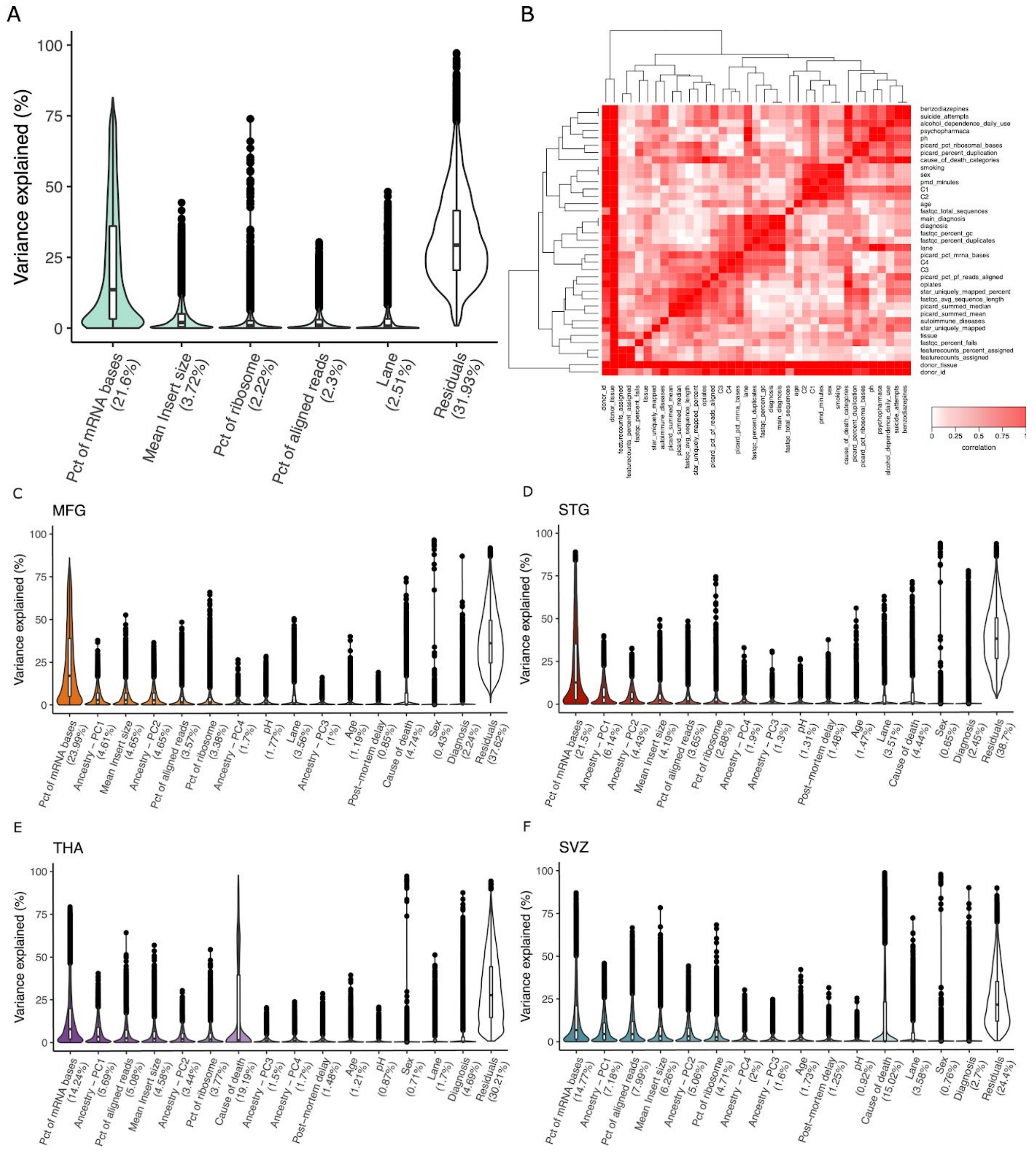


**Supplementary Figure 5: Main sources of expression variation and correlation of covariates.**

A) Percentage of variance explained by technical factors for 255 samples for each gene, variancePartition package (Hoffman and Schadt 2016). Numbers between parenthesis indicate mean values. B) Pairwise canonical correlation between the covariates. Red indicates high correlation; white indicates low correlation. C-F) Percentage of variance explained by technical and biological covariates by each brain region.

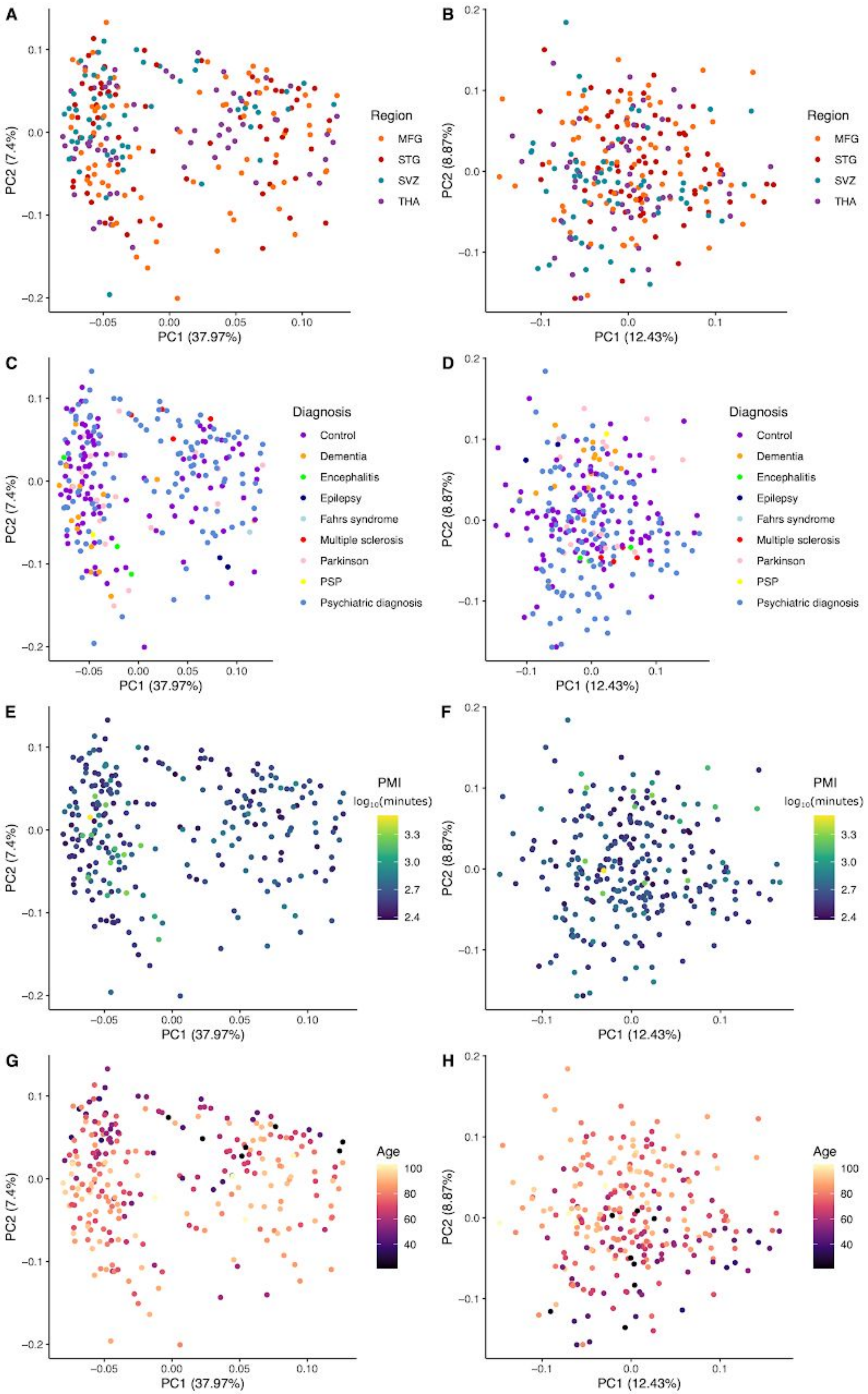

A
B
C
D
E
F
G
H
PC1 (37.97%)
PC2 (7.4%)
PC1 (12.43%)
PC2 (8.87%)
Region
MFG
STG
SVZ
THA
Diagnosis
Control
Dementia
Encephalitis
Epilepsy
Fahrs syndrome
Multiple sclerosis
Parkinson
PSP
Psychiatric diagnosis
PMI
log10(minutes)
Age

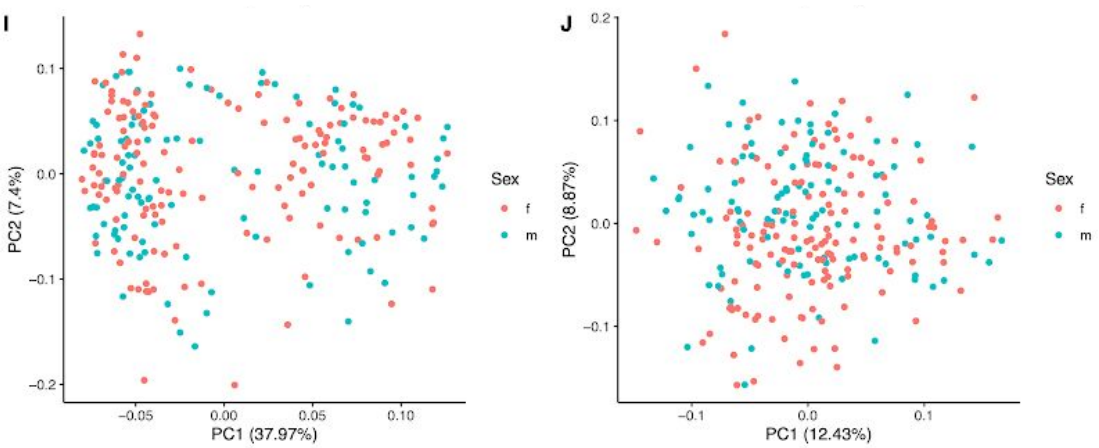


**Supplementary Figure 6: Principal component analysis (PCAs) and data adjustment.**
Pair of PCA before (left A,C,E,G) and after (right B,D,F,H) correction by regressing out technical confounders, colored by A-B) region; C-D) diagnosis; E-F) post-mortem interval (minutes); G-H) age; I-J) sex. The plots show voom-TMM normalized expression for all samples.

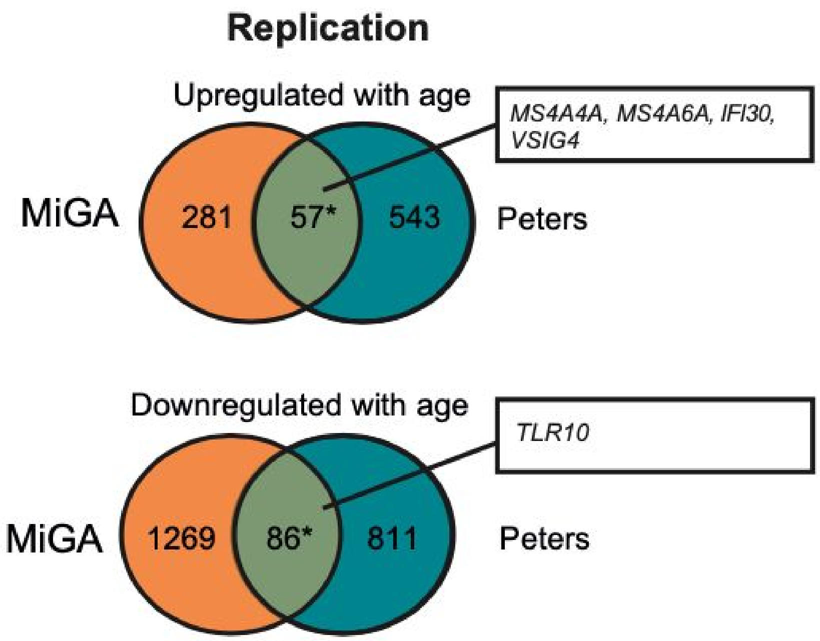


**Supplementary Figure 7: Replication of MiGA age-related genes with peripheral human blood.**

Upregulated in age between MiGA and Peters et al. (Peters et al. 2015). (OR 7.14; 95% CI = 5.21-9.66, $P < 1e-16$). Downregulated in age between MiGA and Peters (OR = 1.49; 95% CI = 1.17-1.88; P = 6e-4).

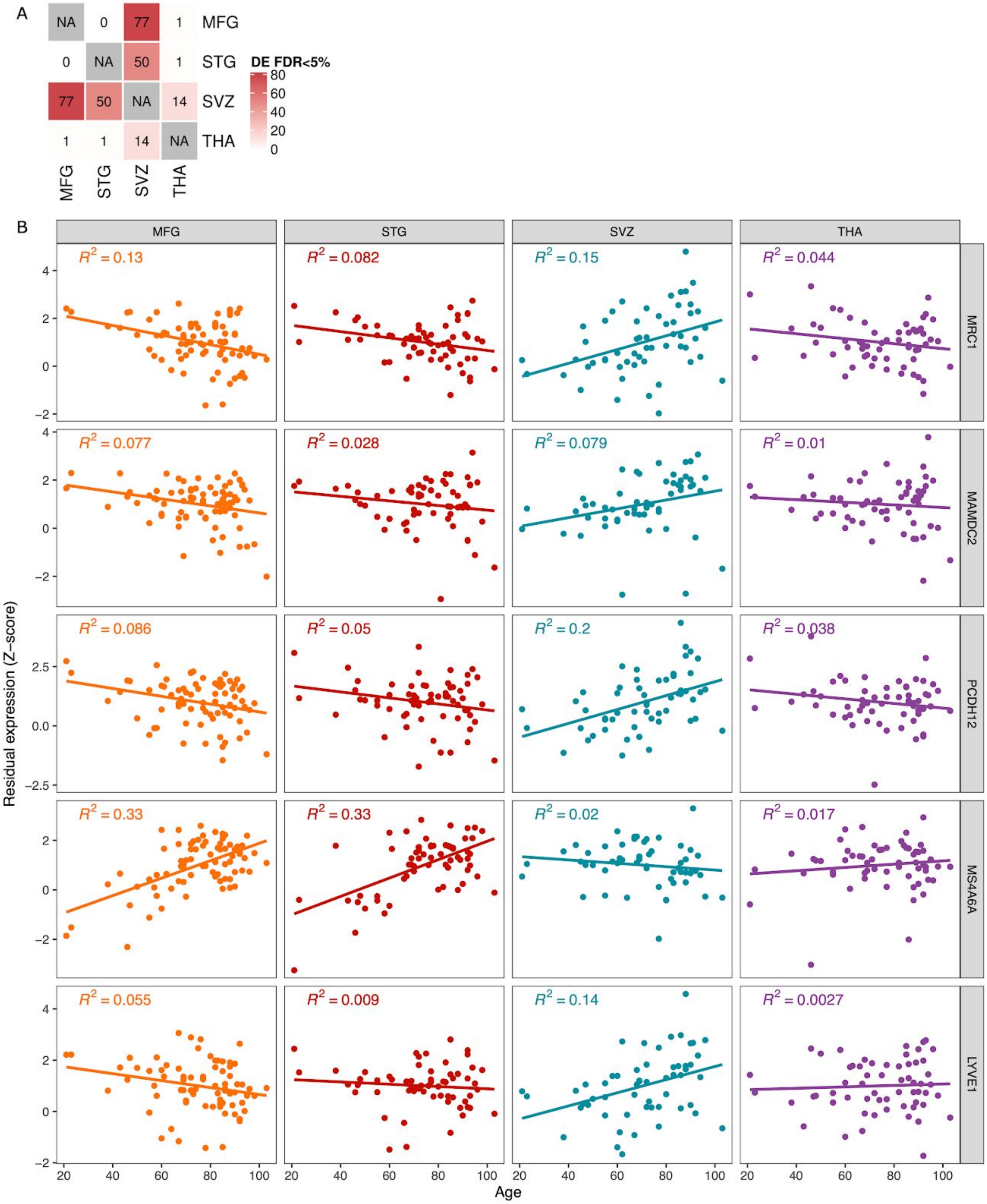


**Supplementary Figure 8: Differential expression by age interaction with brain regions.**

A) Number of pairwise DE genes (FDR<0.05) using a linear model with interaction between age and region. B) Top five genes prioritized by the DE analysis to show flipped effects in direction of gene expression per age and region.

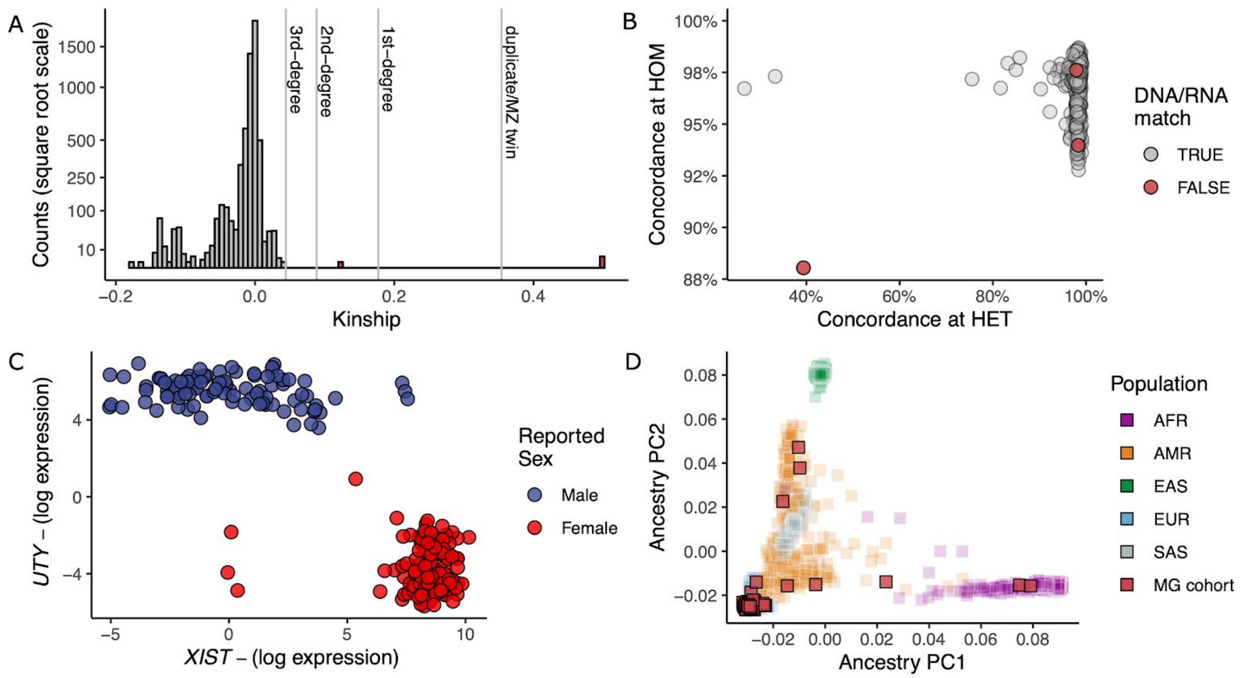


**Supplementary Figure 9: Genotyping QC.**

A) Distribution of number of samples per estimated kinship measured using KING. Estimated kinship coefficient ranges of >0.354, [0.177, 0.354], [0.0884, 0.177] and [0.0442, 0.0884] corresponds to duplicate or monozygotic twins, 1st-degree, 2nd-degree, and 3rd-degree relationships respectively. B) DNA-RNA sample matching using QTLtools-mbv measured by the percentage of concordance at heterozygous and homozygous genotypes. Samples in red failed to match ids between each data. C) Log scaled expression (*voom*) of sex chromosome genes (*UTY* and *XIST*) for each donor colored by reported sex (blue = male, red = female). D) First two ancestry principal components of MiGA samples (red squares) on top of 1000 Genome individuals colored by major populations. African [AFR], Admixed American [AMR], East Asian [EAS], European [EUR], South Asian [SAS].

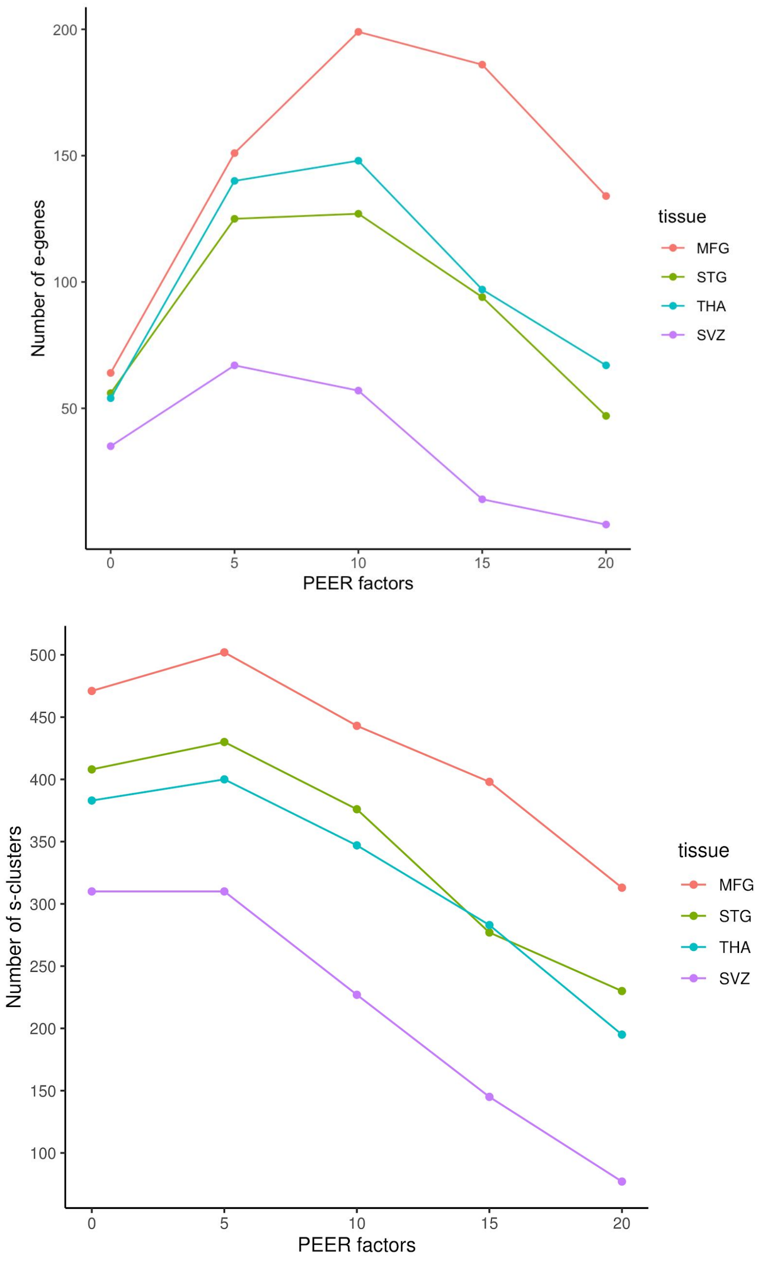


**Supplementary Figure 10: Probabilistic Estimation of Expression Residuals (PEER) correction for non-genetic factors in eQTL and sQTL analyses.**

To account for hidden effects in gene expression data, such as technical artifacts, we used PEER. After adjusting for age and sex, a number of PEER factors (from 0 to 20) was tested to maximize eQTLs per gene (top) and splicing QTLs per intron cluster (bottom) discovery at q-val < 0.05. Based on this result, 5 (SVZ) and 10 (MFG, STG, and THA) PEER factors were selected for eQTLs and 0 (SVZ) and 5 (MFG, STG, and THA) PEER factors for sQTLs. Factors were regressed out of the respective gene expression or splicing matrix before running the association tests.

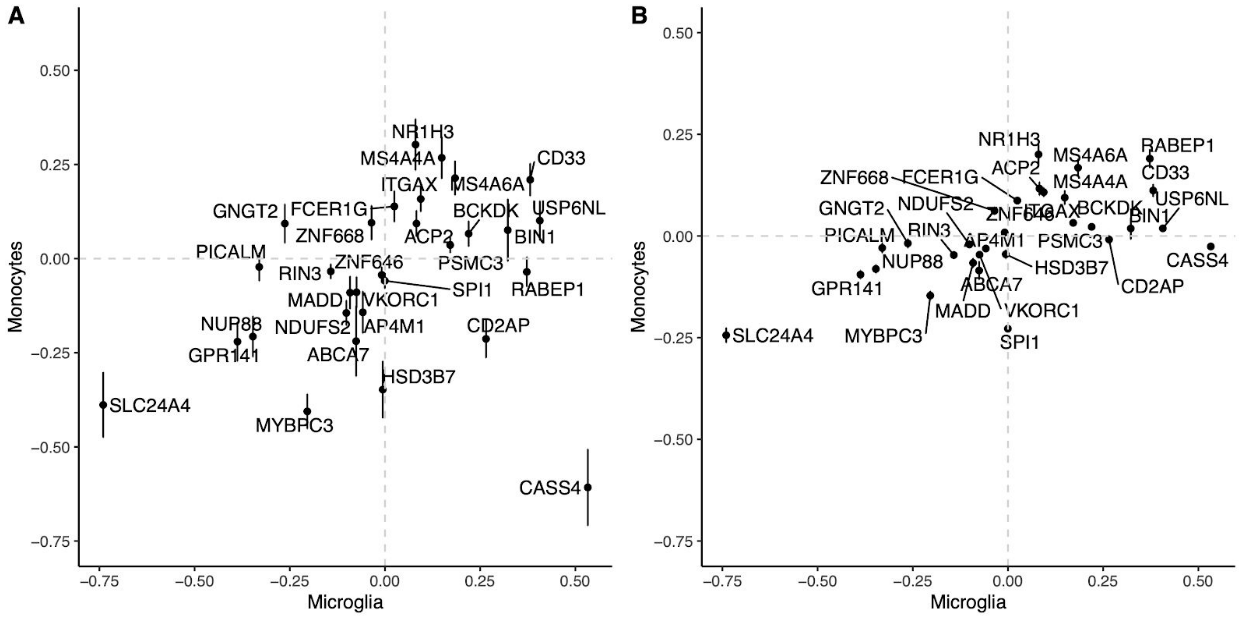


**Supplementary Figure 11: eQTL effect sizes of the AD-associated genes.**

A) x axis shows the eQTL betas from microglia dataset from this study (MiGA), and y axis shows the eQTL betas of monocytes from MyND dataset (Navarro et al.). B) x axis shows the eQTL betas from microglia (MiGA), and y axis shows the eQTL betas of monocytes from Fairfax et al. Vertical lines indicate standard error.

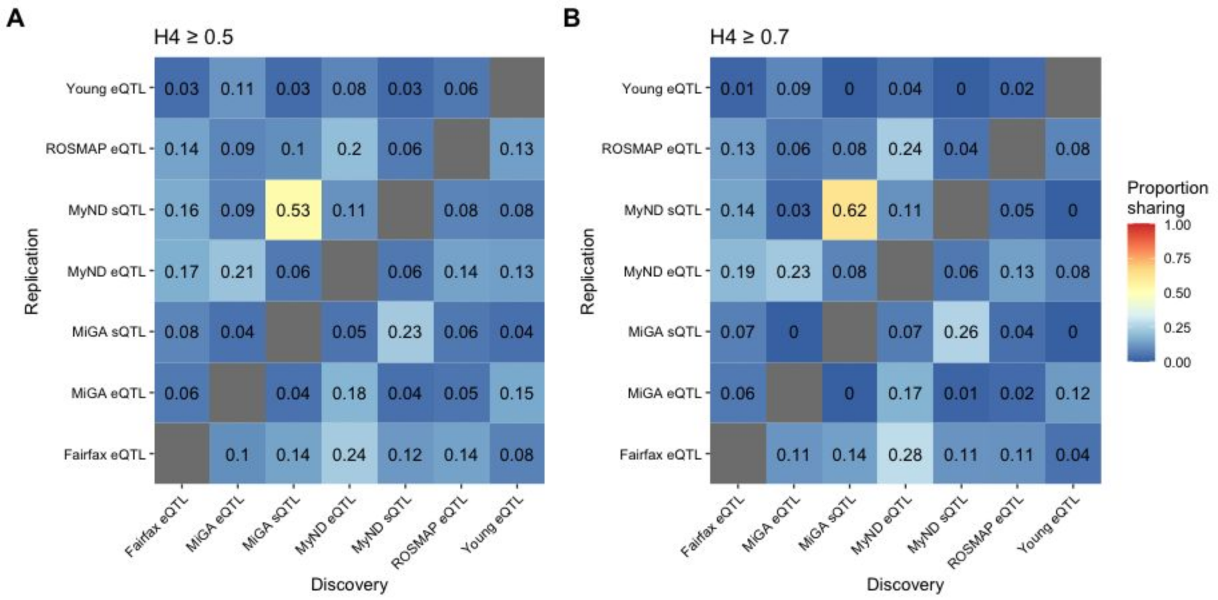


**Supplementary Figure 12: Pairwise sharing of colocalized genes.**

Locus-Gene combinations across all 5 diseases that colocalized at PP4 >= 0.5 (**A**) or >= 0.7 (**B**) were compared between pairs of QTL datasets. For example, 53% of genes that colocalised PP4 > 0.5 in MiGA sQTLs also colocalized in MyND sQTLs, whereas only 23% of the MyND sQTLs were found in the MiGA sQTLs.

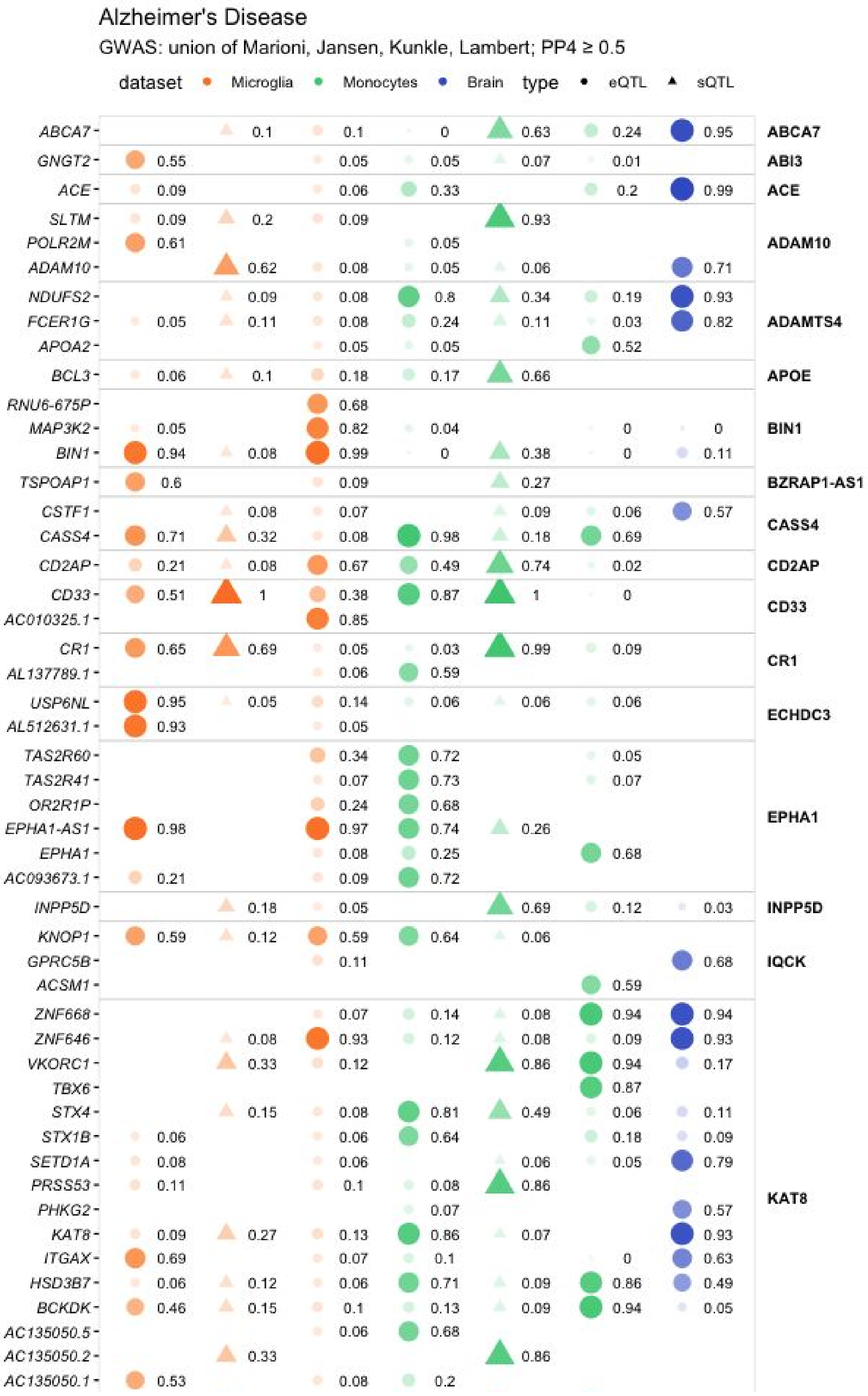
Alzheimer's Disease
GWAS: union of Marioni, Jansen, Kunkle, Lambert; PP4 ≥ 0.5
dataset Microglia Monocytes Brain type eQTL sQTL
ABCA7
GNGT2
ACE
SLTM
POLR2M
ADAM10
NDUFS2
FCER1G
APOA2
BCL3
RNU6-675P
MAP3K2
BIN1
TSPOAP1
CSTF1
CASS4
CD2AP
CD33
AC010325.1
CR1
AL137789.1
USP6NL
AL512631.1
TAS2R60
TAS2R41
OR2R1P
EPHA1-AS1
EPHA1
AC093673.1
INPP5D
KNOP1
GPRC5B
ACSM1
ZNF668
ZNF646
VKORC1
TBX6
STX4
STX1B
SETD1A
PRSS53
PHKG2
KAT8
ITGAX
HSD3B7
BCKDK
AC135050.5
AC135050.2
AC135050.1
ABCA7
ABI3
ACE
ADAM10
ADAMTS4
APOE
BIN1
BZRAP1-AS1
CASS4
CD2AP
CD33
CR1
ECHDC3
EPHA1
INPP5D
IQCK
KAT8

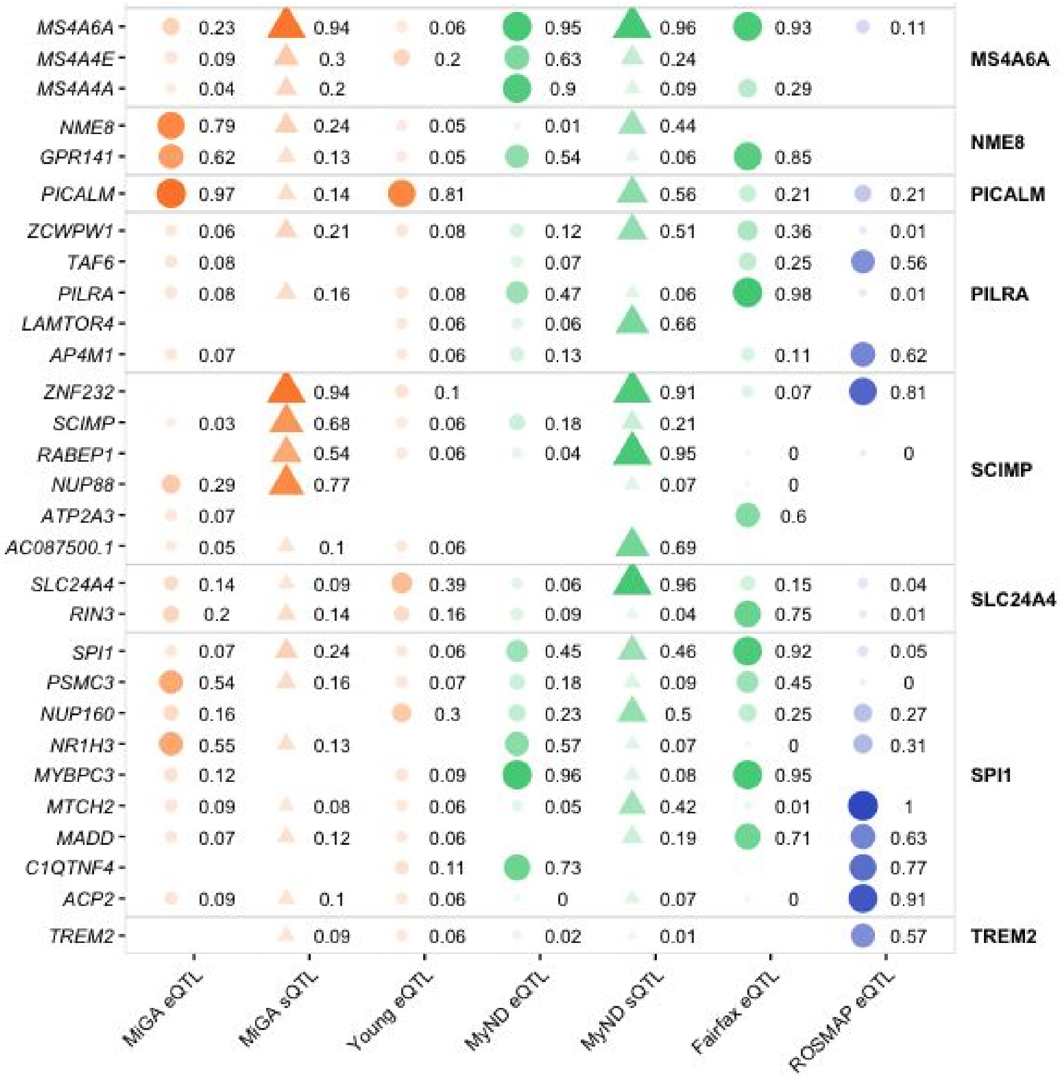


**Supplementary Figure 13: Full colocalization results in Alzheimer's Disease.**

Colocalization PP4 displayed for each GWAS locus (right text) and gene (left text) for each QTL dataset. An empty value means no QTL was present for testing for that gene in that dataset.

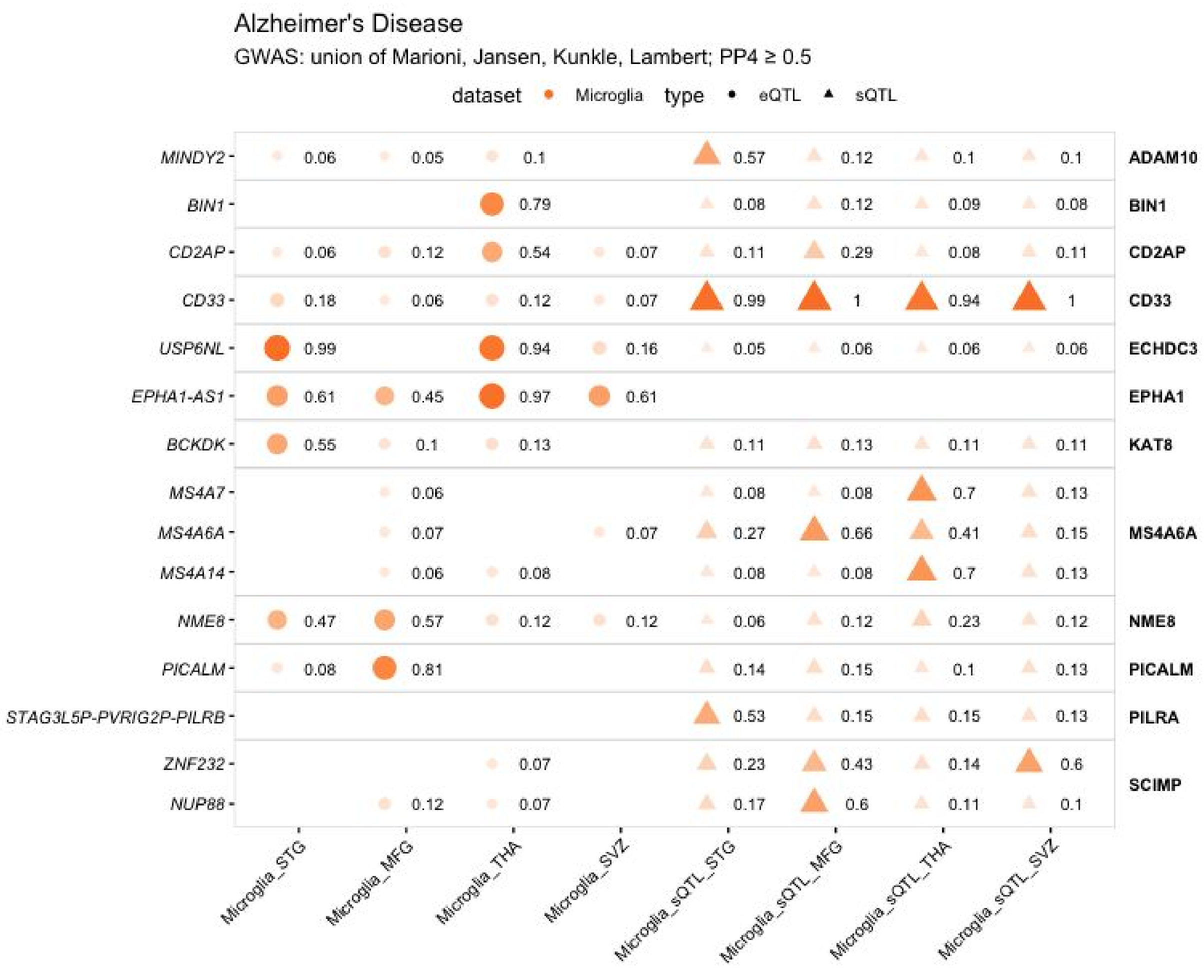


**Supplementary Figure 14: Colocalization results for each regional microglia dataset in Alzheimer's Disease.**

Colocalization PP4 displayed for each GWAS locus (right text) and gene (left text) for each QTL dataset. An empty value means no QTL was present for testing for that gene in that dataset.

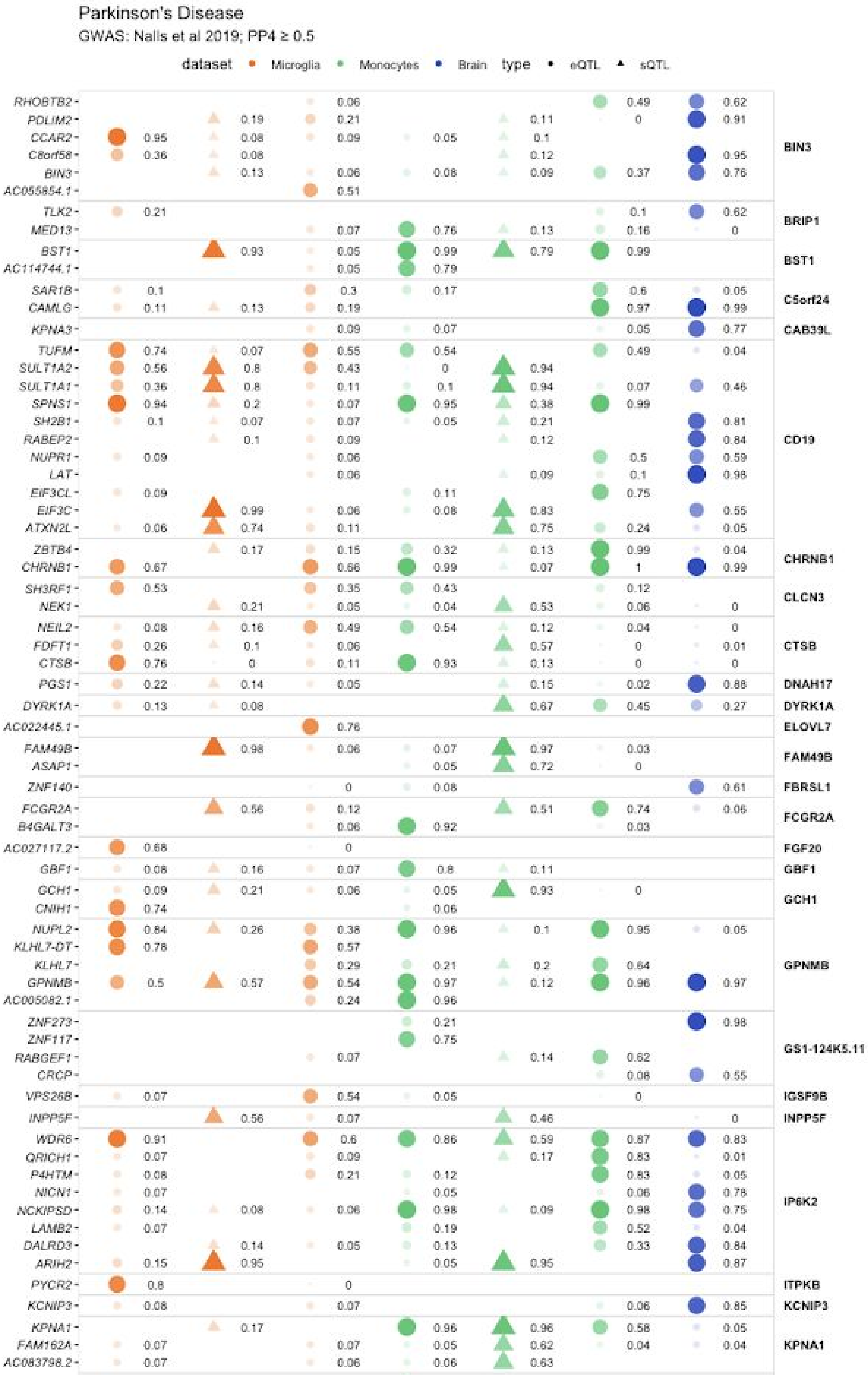
Parkinson's Disease
GWAS: Nalls et al 2019; PP4 ≥ 0.5
dataset
Microglia
Monocytes
Brain
type
eQTL
sQTL
RHOBTB2
PDLIM2
CCAR2
C8orf58
BIN3
AC055854.1
TLK2
MED13
BST1
AC114744.1
SAR1B
CAMLG
KPNA3
TUFM
SULT1A2
SULT1A1
SPNS1
SH2B1
RABEP2
NUPR1
LAT
EIF3CL
EIF3C
ATXN2L
ZBTB4
CHRNB1
SH3RF1
NEK1
NEIL2
FDFT1
CTSB
PGS1
DYRK1A
AC022445.1
FAM49B
ASAP1
ZNF140
FCGR2A
B4GALT3
AC027117.2
GBF1
GCH1
CNIH1
NUPL2
KLHL7-DT
KLHL7
GPNMB
AC005082.1
ZNF273
ZNF117
RABGEF1
CRCP
VPS26B
INPP5F
WDR6
QRICH1
P4HTM
NICN1
NCKIPSD
LAMB2
DALRD3
ARIH2
PYCR2
KCNIP3
KPNA1
FAM162A
AC083798.2
BIN3
BRIP1
BST1
C5orf24
CAB39L
CD19
CHRNB1
CLCN3
CTSB
DNAH17
DYRK1A
ELOVL7
FAM49B
FBRSL1
FCGR2A
FGF20
GBF1
GCH1
GPNMB
GS1-124K5.11
IGSF9B
INPP5F
IP6K2
ITPKB
KCNIP3
KPNA1

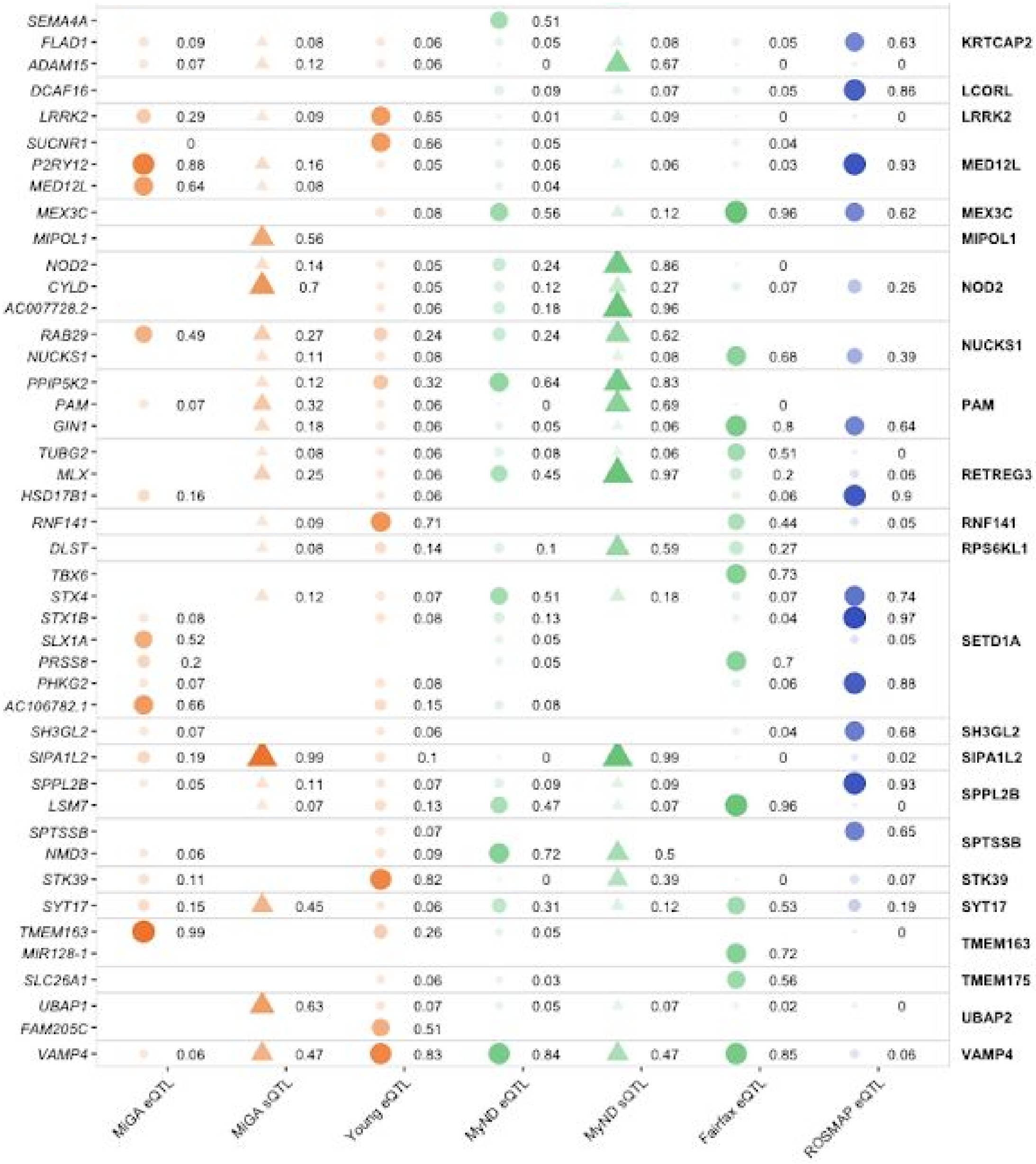

**Supplementary Figure 15: Full colocalization results in Parkinson's Disease.**
Colocalization PP4 displayed for each GWAS locus (right text) and gene (left text) for each QTL dataset. An empty value means no QTL was present for testing for that gene in that dataset.

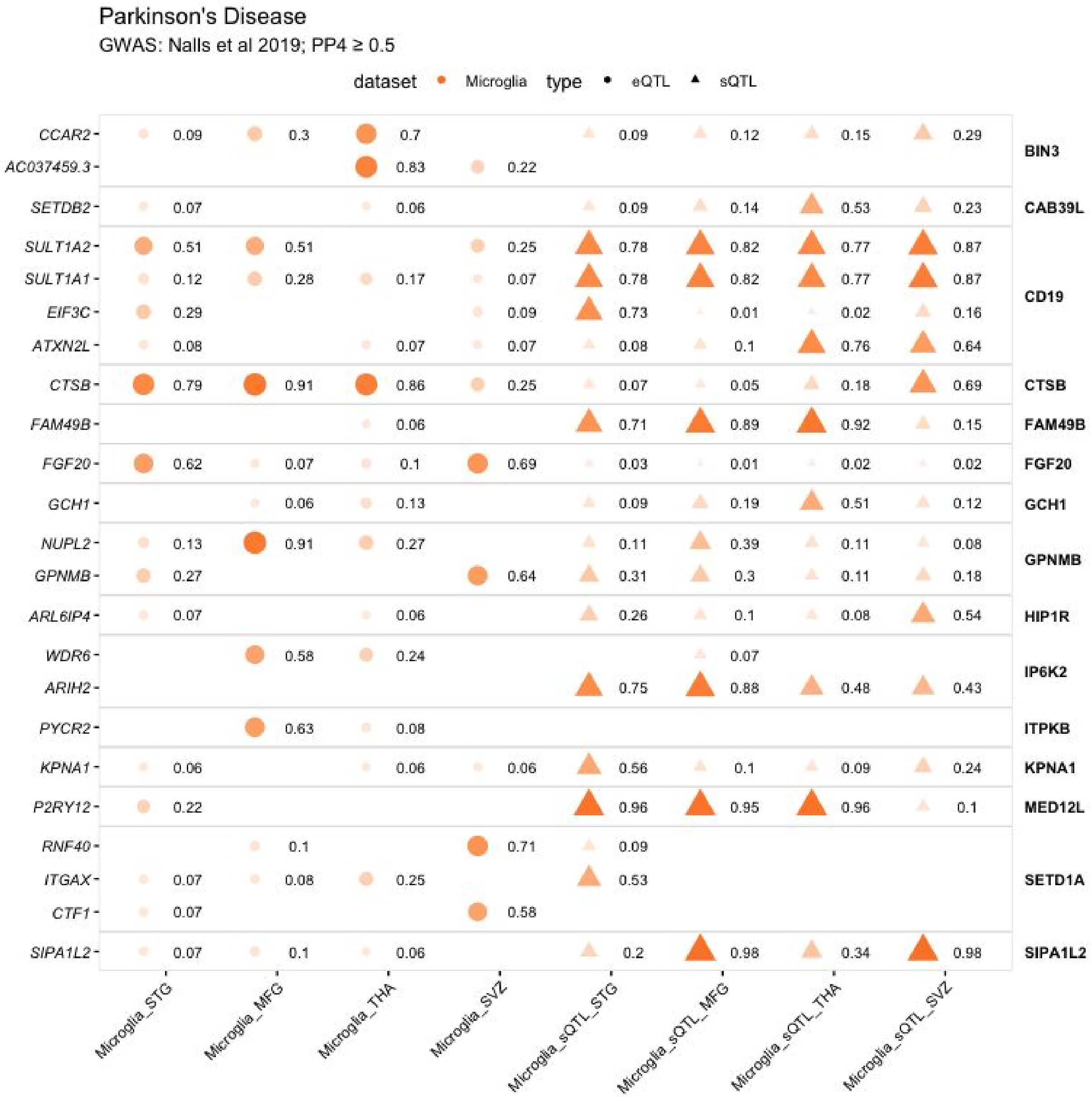


**Supplementary Figure 16: Colocalization results for each regional microglia dataset in Parkinson's Disease.**

Colocalization PP4 displayed for each GWAS locus (right text) and gene (left text) for each QTL dataset. An empty value means no QTL was present for testing for that gene in that dataset.

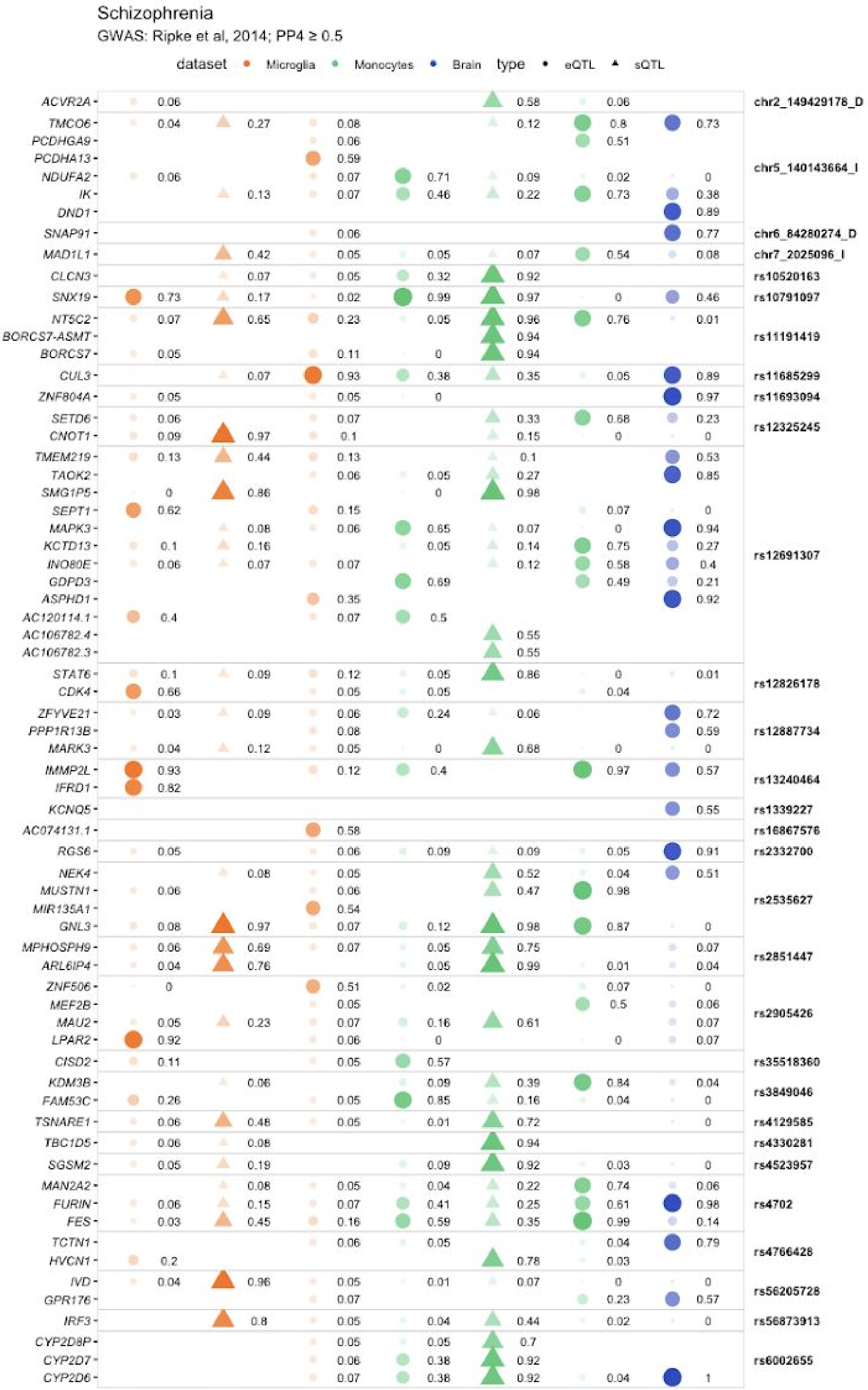
Schizophrenia
GWAS: Ripke et al, 2014; PP4 ≥ 0.5
dataset Microglia Monocytes Brain type eQTL sQTL
ACVR2A 0.06 0.58 0.06 chr2_149429178_D
TMCO6 0.04 0.27 0.08 0.12 0.8 0.73
PCDHGA9 0.06 0.51
PCDHA13 0.59
NDUFA2 0.06 0.07 0.71 0.09 0.02 0
IK 0.13 0.07 0.46 0.22 0.73 0.38
DND1 0.89
chr5_140143664_I
SNAP91 0.06 0.77 chr6_84280274_D
MAD1L1 0.42 0.05 0.05 0.07 0.54 0.08 chr7_2025096_I
CLCN3 0.07 0.05 0.32 0.92 rs10520163
SNX19 0.73 0.17 0.02 0.99 0.97 0 0.46 rs10791097
NT5C2 0.07 0.65 0.23 0.05 0.96 0.76 0.01
BORCS7-ASMT 0.94
BORCS7 0.05 0.11 0 0.94
rs11191419
CUL3 0.07 0.93 0.38 0.35 0.05 0.89 rs11685299
ZNF804A 0.05 0.05 0 0.97 rs11693094
SETD6 0.06 0.07 0.33 0.68 0.23
CNOT1 0.09 0.97 0.1 0.15 0 0
rs12325245
TMEM219 0.13 0.44 0.13 0.1 0.53
TAOK2 0.06 0.05 0.27 0.85
SMG1P5 0 0.86 0 0.98
SEPT1 0.62 0.15 0.07 0
MAPK3 0.08 0.06 0.65 0.07 0 0.94
KCTD13 0.1 0.16 0.05 0.14 0.75 0.27
INO80E 0.06 0.07 0.07 0.12 0.58 0.4
GDPD3 0.69 0.49 0.21
ASPHD1 0.35 0.92
AC120114.1 0.4 0.07 0.5
AC106782.4 0.55
AC106782.3 0.55
rs12691307
STAT6 0.1 0.09 0.12 0.05 0.86 0 0.01
CDK4 0.66 0.05 0.05 0.04
rs12826178
ZFYVE21 0.03 0.09 0.06 0.24 0.06 0.72
PPP1R13B 0.08 0.59
MARK3 0.04 0.12 0.05 0 0.68 0 0
rs12887734
IMMP2L 0.93 0.12 0.4 0.97 0.57
IFRD1 0.82
rs13240464
KCNQ5 0.55 rs1339227
AC074131.1 0.58 rs16867576
RGS6 0.05 0.06 0.09 0.09 0.05 0.91 rs2332700
NEK4 0.08 0.05 0.52 0.04 0.51
MUSTN1 0.06 0.06 0.47 0.98
MIR135A1 0.54
GNL3 0.08 0.97 0.07 0.12 0.98 0.87 0
rs2535627
MPHOSPH9 0.06 0.69 0.07 0.05 0.75 0.07
ARL6IP4 0.04 0.76 0.05 0.99 0.01 0.04
rs2851447
ZNF506 0 0.51 0.02 0.07 0
MEF2B 0.05 0.5 0.06
MAU2 0.05 0.23 0.07 0.16 0.61 0.07
LPAR2 0.92 0.06 0 0 0.07
rs2905426
CISD2 0.11 0.05 0.57 rs35518360
KDM3B 0.06 0.09 0.39 0.84 0.04
FAM53C 0.26 0.05 0.85 0.16 0.04 0
rs3849046
TSNARE1 0.06 0.48 0.05 0.01 0.72 0 rs4129585
TBC1D5 0.06 0.08 0.94 rs4330281
SGSM2 0.05 0.19 0.09 0.92 0.03 0 rs4523957
MAN2A2 0.08 0.05 0.04 0.22 0.74 0.06
FURIN 0.06 0.15 0.07 0.41 0.25 0.61 0.98
FES 0.03 0.45 0.16 0.59 0.35 0.99 0.14
rs4702
TCTN1 0.06 0.05 0.04 0.79
HVCN1 0.2 0.78 0.03
rs4766428
IVD 0.04 0.96 0.05 0.01 0.07 0 0
GPR176 0.07 0.23 0.57
rs56205728
IRF3 0.8 0.05 0.04 0.44 0.02 0 rs56873913
CYP2D8P 0.05 0.05 0.7
CYP2D7 0.06 0.38 0.92
CYP2D6 0.07 0.38 0.92 0.04 1
rs6002655

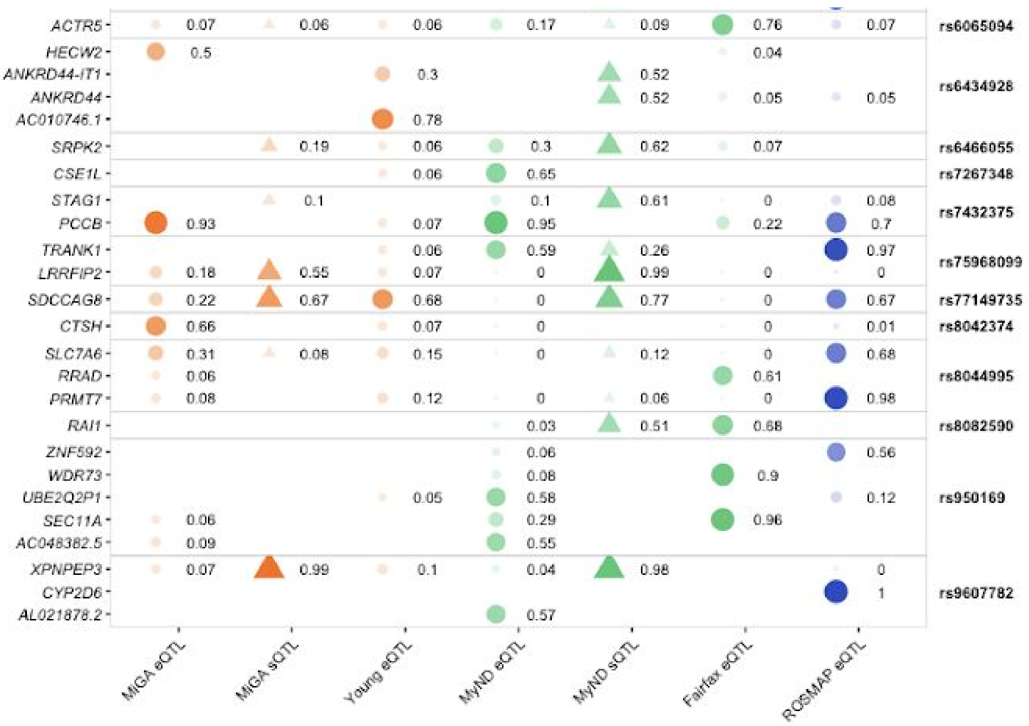


**Supplementary Figure 17: Full colocalization results in schizophrenia.**
Colocalization PP4 displayed for each GWAS locus (right text) and gene (left text) for each QTL dataset. An empty value means no QTL was present for testing for that gene in that dataset.

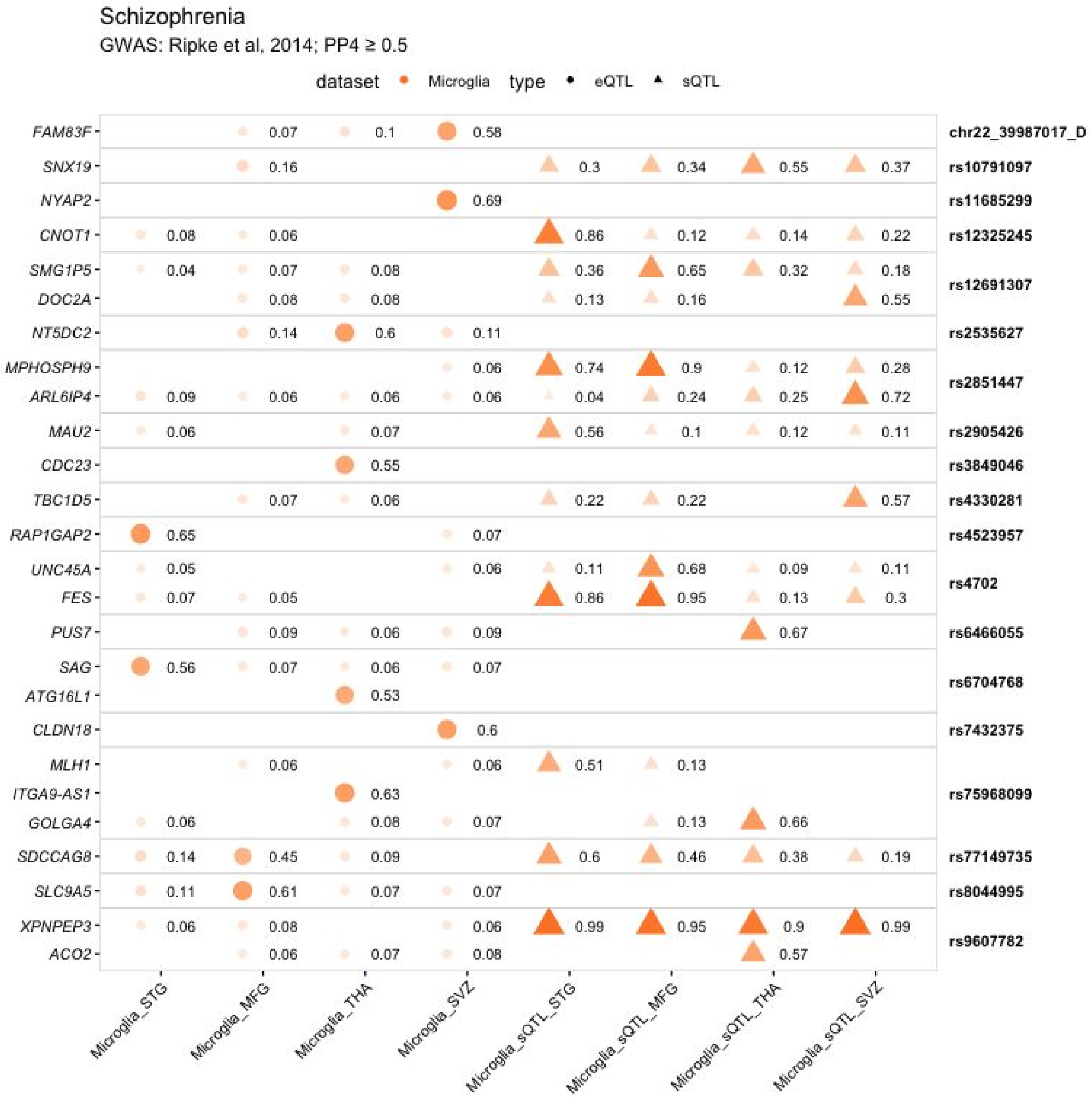


**Supplementary Figure 18: Colocalization results for each regional microglia dataset in schizophrenia.**

Colocalization PP4 displayed for each GWAS locus (right text) and gene (left text) for each QTL dataset. An empty value means no QTL was present for testing for that gene in that dataset.

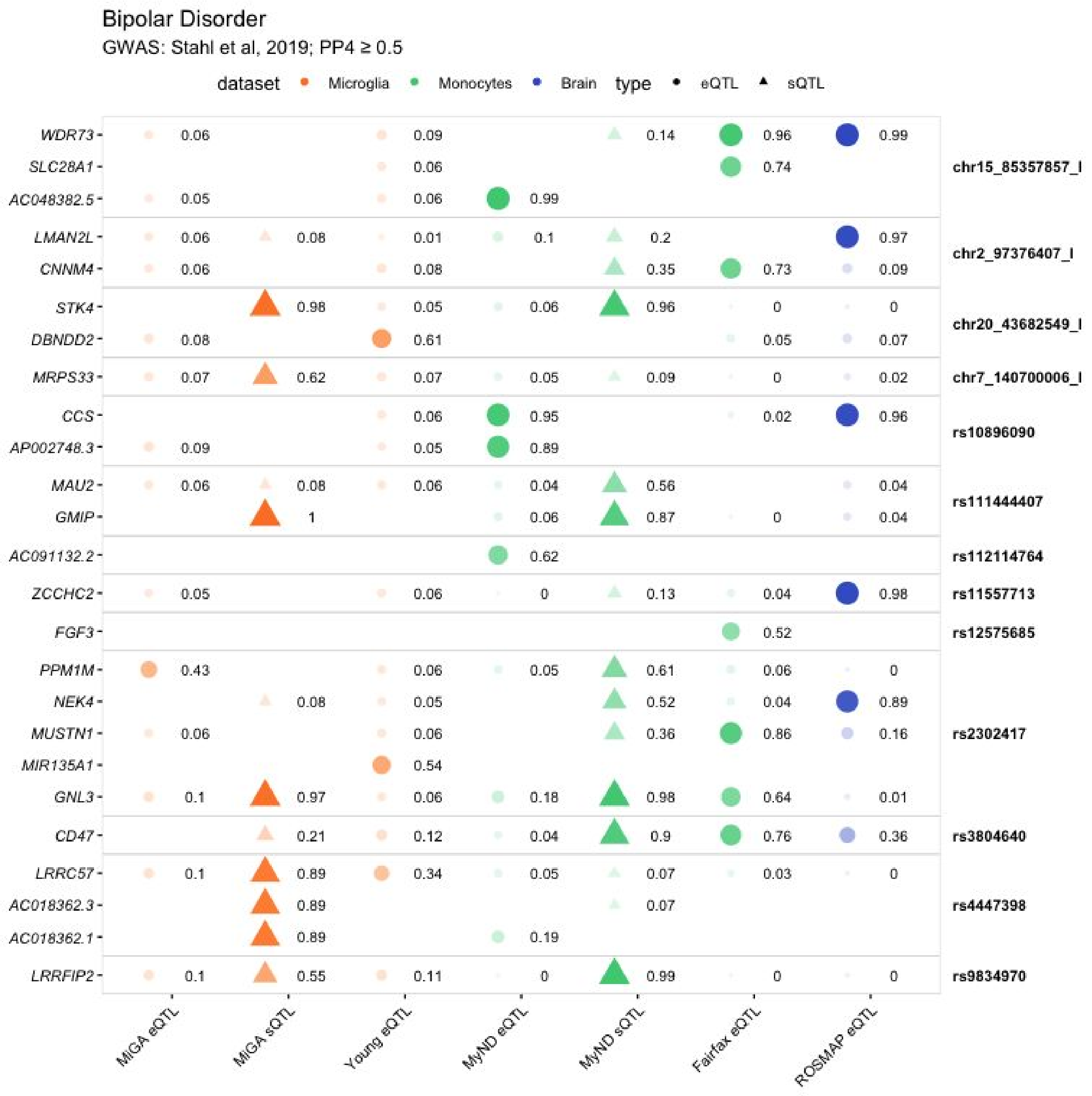


**Supplementary Figure 19: Full colocalization results in bipolar disorder.**
Colocalization PP4 displayed for each GWAS locus (right text) and gene (left text) for each QTL dataset. An empty value means no QTL was present for testing for that gene in that dataset.

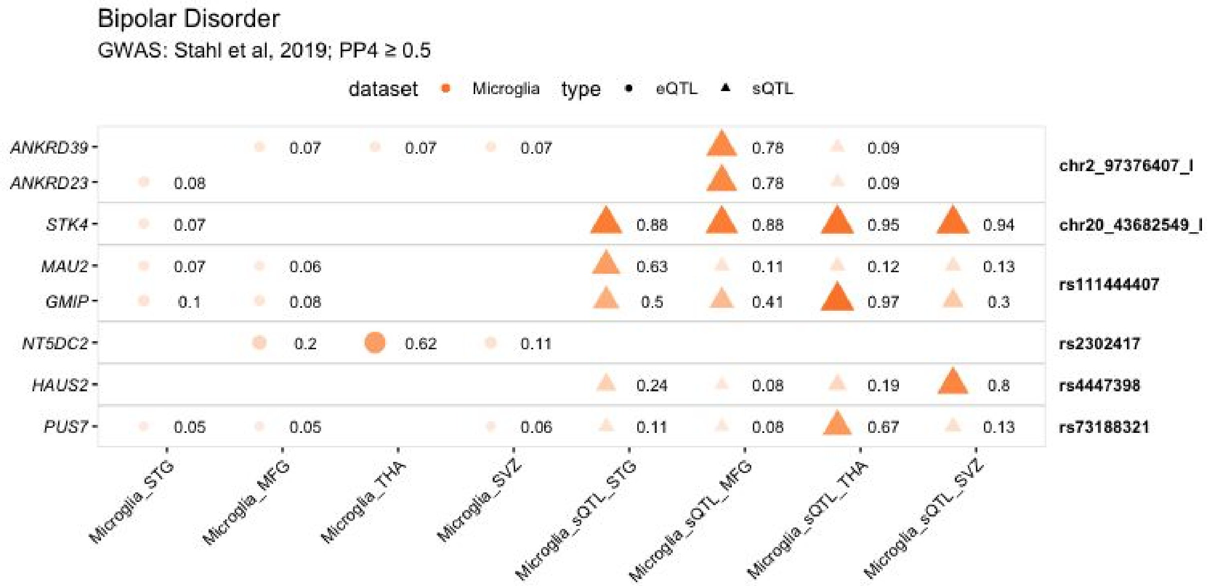


**Supplementary Figure 20: Colocalization results for each regional microglia dataset in bipolar disorder.**

Colocalization PP4 displayed for each GWAS locus (right text) and gene (left text) for each QTL dataset. An empty value means no QTL was present for testing for that gene in that dataset.

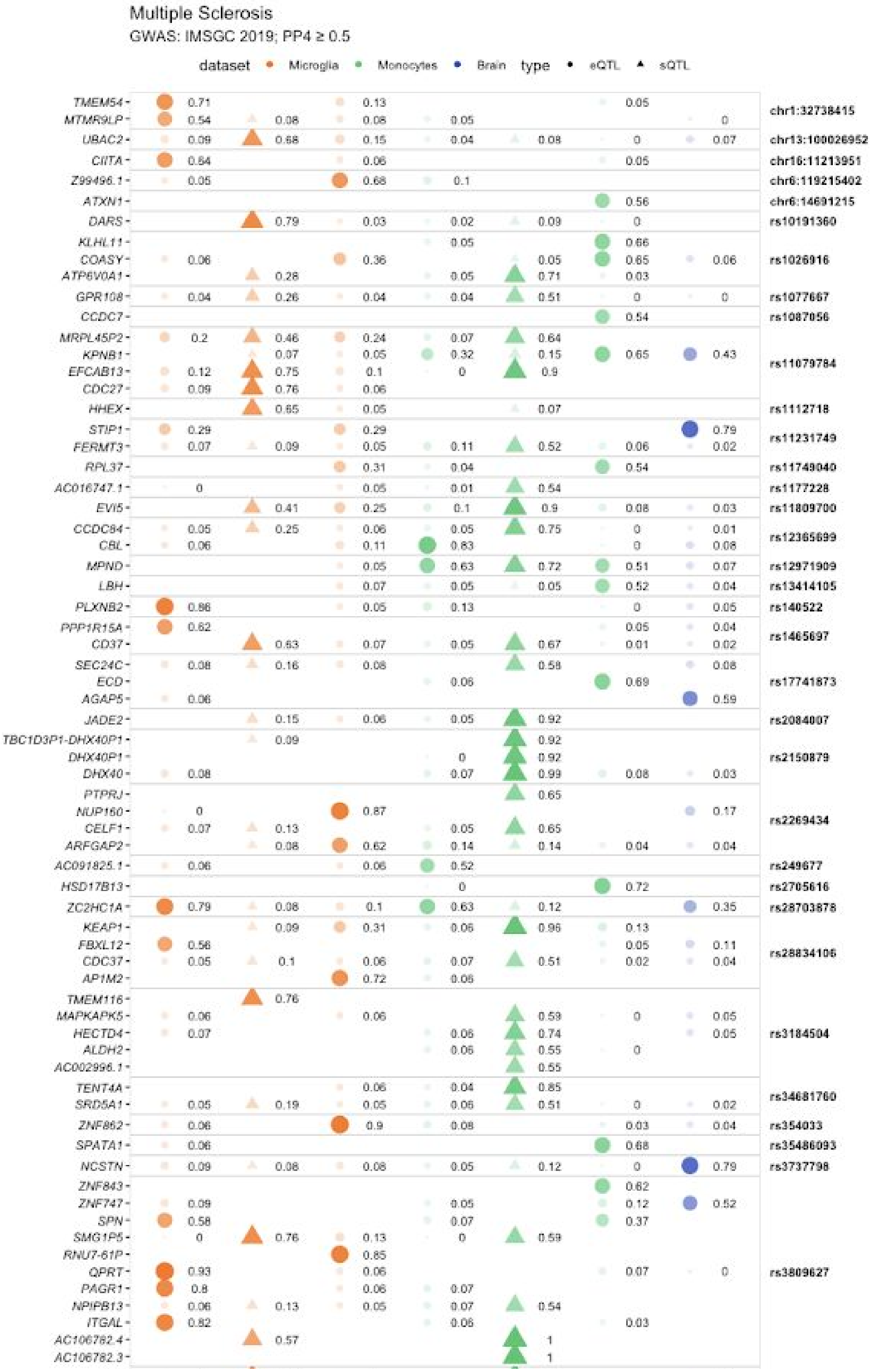
Multiple Sclerosis
GWAS: IMSGC 2019; PP4 ≥ 0.5
dataset Microglia Monocytes Brain type eQTL sQTL
TMEM54
MTMR9LP
UBAC2
CIITA
Z99496.1
ATXN1
DARS
KLHL11
COASY
ATP6V0A1
GPR108
CCDC7
MRPL45P2
KPNB1
EFCAB13
CDC27
HHEX
STIP1
FERMT3
RPL37
AC016747.1
EVI5
CCDC84
CBL
MPND
LBH
PLXNB2
PPP1R15A
CD37
SEC24C
ECD
AGAP5
JADE2
TBC1D3P1-DHX40P1
DHX40P1
DHX40
PTPRJ
NUP160
CELF1
ARFGAP2
AC091825.1
HSD17B13
ZC2HC1A
KEAP1
FBXL12
CDC37
AP1M2
TMEM116
MAPKAPK5
HECTD4
ALDH2
AC002996.1
TENT4A
SRD5A1
ZNF862
SPATA1
NCSTN
ZNF843
ZNF747
SPN
SMG1P5
RNU7-61P
QPRT
PAGR1
NPIPB13
ITGAL
AC106782.4
AC106782.3
chr1:32738415
chr13:100026952
chr16:11213951
chr6:119215402
chr6:14691215
rs10191360
rs1026916
rs1077667
rs1087056
rs11079784
rs1112718
rs11231749
rs11749040
rs1177228
rs11809700
rs12365699
rs12971909
rs13414105
rs140522
rs1465697
rs17741873
rs2084007
rs2150879
rs2269434
rs249677
rs2705616
rs28703878
rs28834106
rs3184504
rs34681760
rs354033
rs35486093
rs3737798
rs3809627

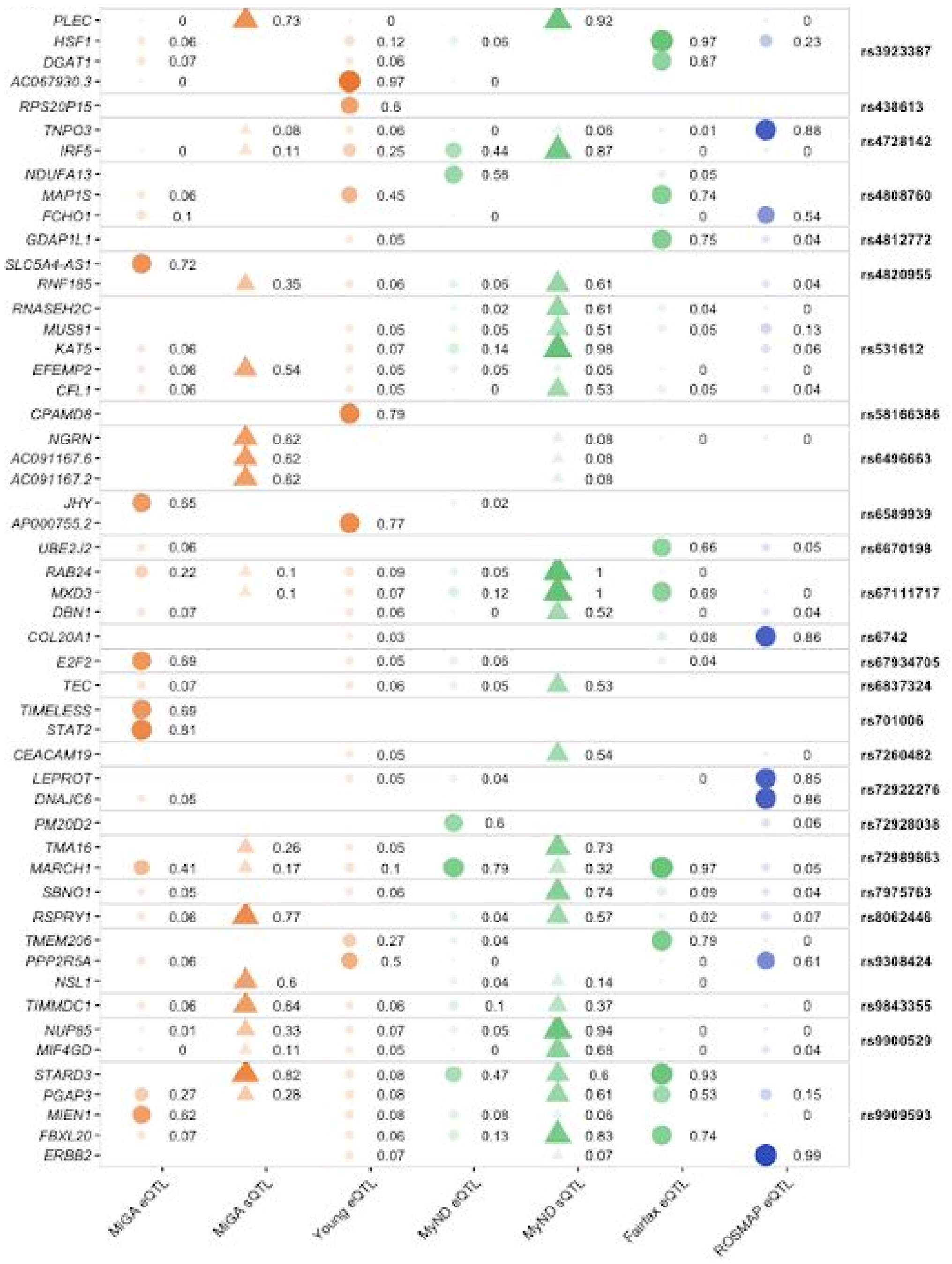


**Supplementary Figure 21: Full colocalization results in multiple sclerosis.**
Colocalization PP4 displayed for each GWAS locus (right text) and gene (left text) for each QTL dataset. An empty value means no QTL was present for testing for that gene in that dataset.

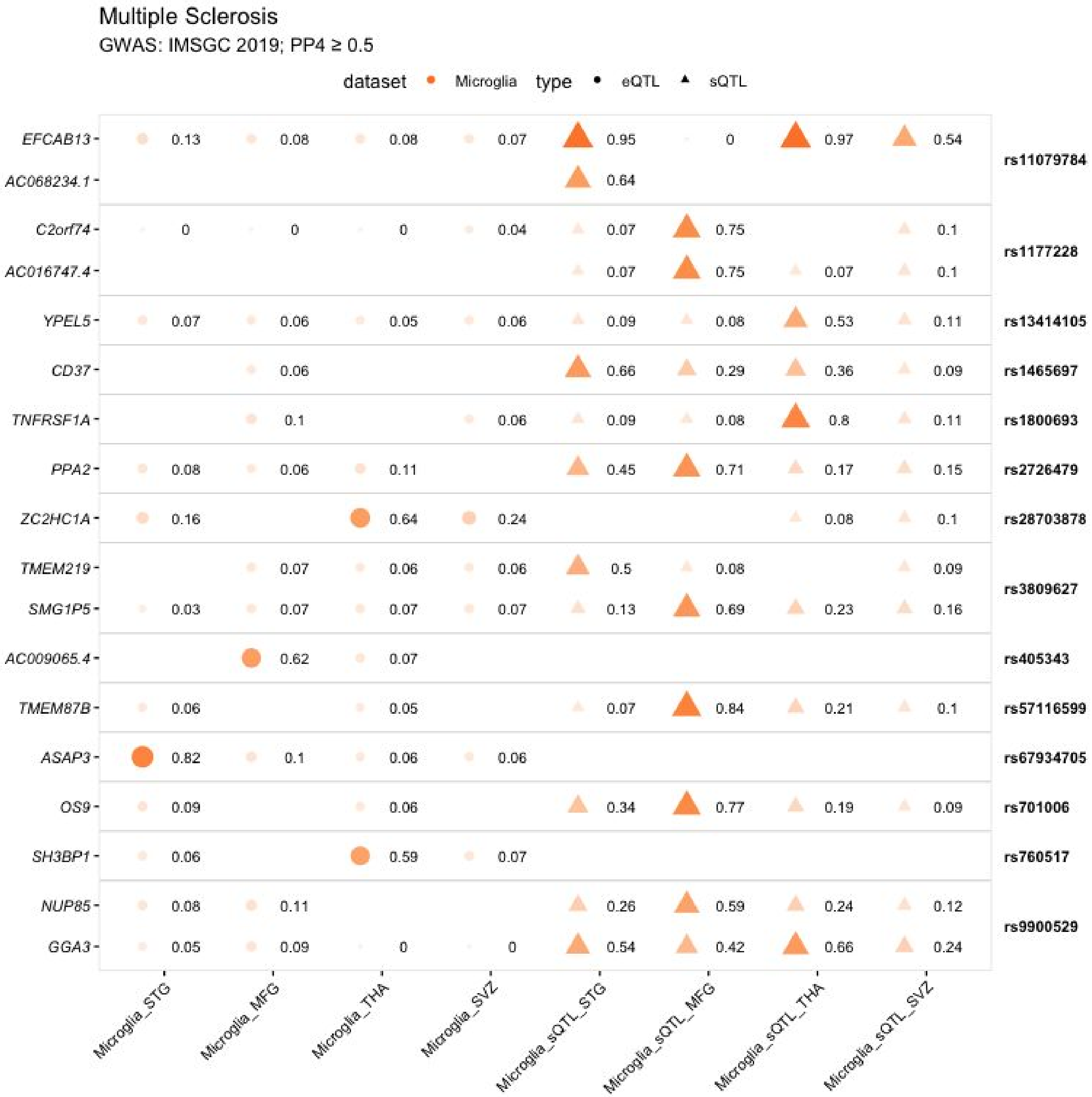


**Supplementary Figure 22: Colocalization results for each regional microglia dataset in multiple sclerosis.**

Colocalization PP4 displayed for each GWAS locus (right text) and gene (left text) for each QTL dataset. An empty value means no QTL was present for testing for that gene in that dataset.

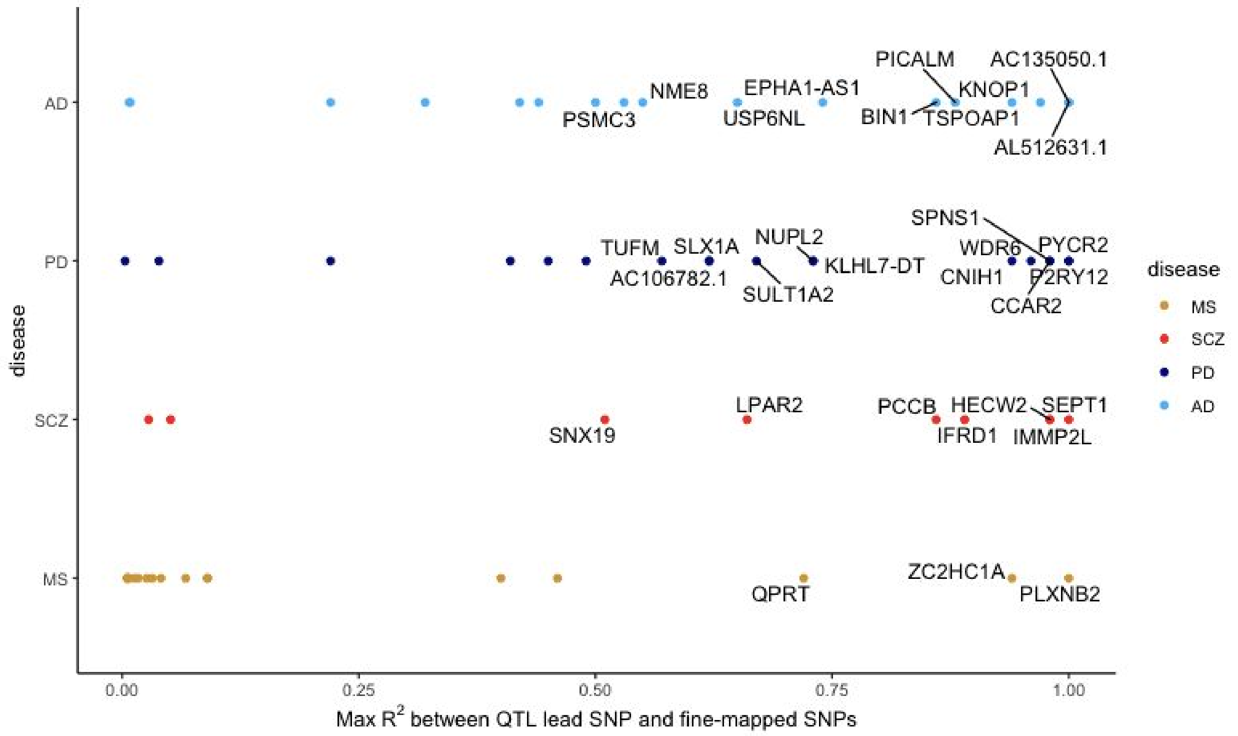


**Supplementary Figure 23: Fine-mapping of loci colocalizing with MiGA eQTLs.**

Genes in each disease plotted by the maximum $R^2$ between the lead QTL SNP and the set of fine-mapped SNPs at that GWAS locus (1000 Genomes phase 3 European samples). Genes with $R^2$ > 0.5 are labelled.

Locus: CD19 GWAS: Nalls et al, 2019

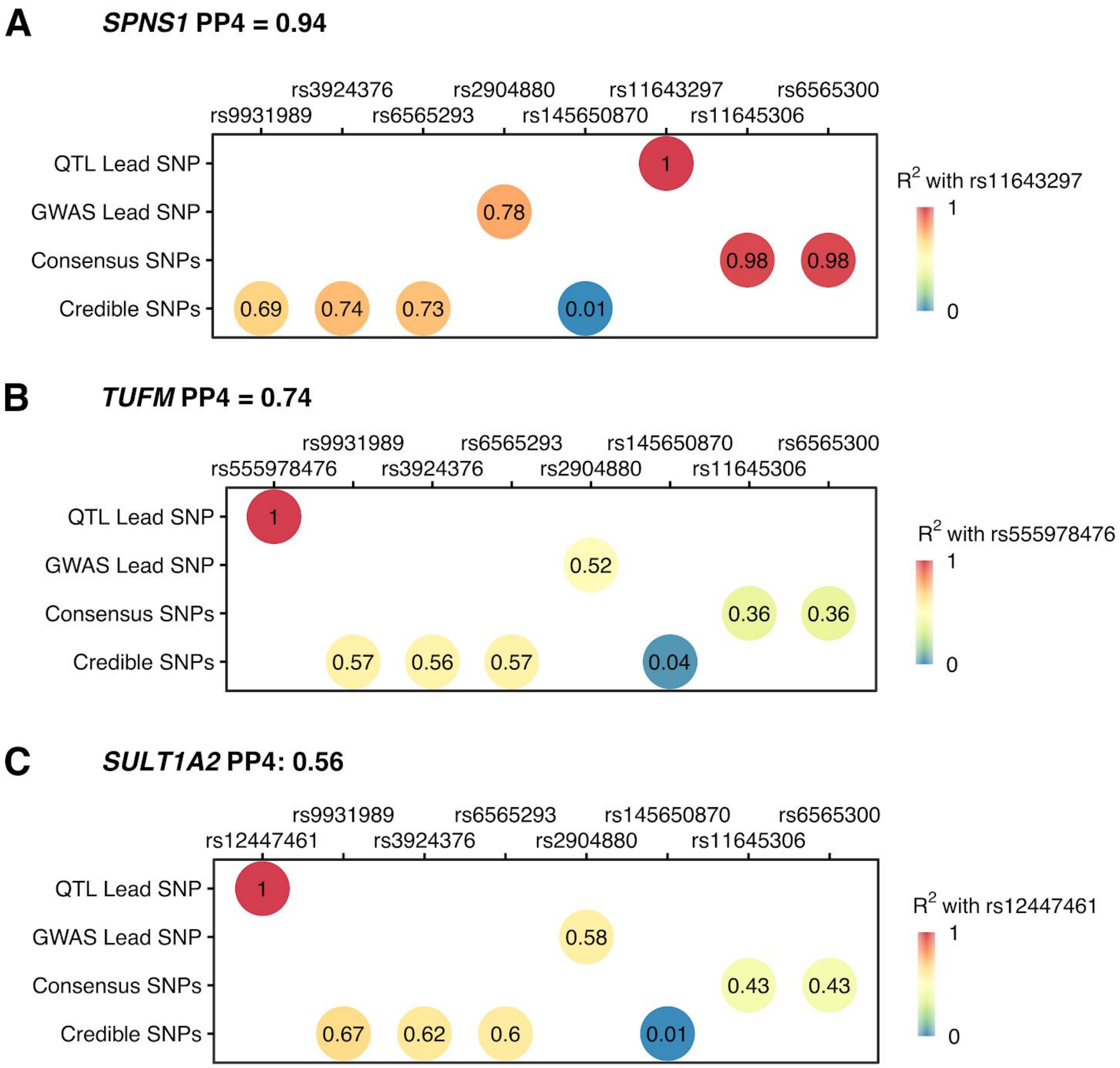


**Supplementary Figure 24: Fine-mapping and LD calculation aids gene prioritization at the CD19 Parkinson's Disease locus.**

The CD19 locus colocalizes with MiGA eQTLs in *SPNS1* (**A**), *TUFM* (**B**), and *SULT1A2* (**C**). Multiple tool fine-mapping with echolocatoR produced two consensus SNPs and four credible SNPs at the CD19 locus. Here we plot the QTL lead SNP, the lead GWAS SNP and the fine-mapped SNPs, coloured and labelled by the LD with the lead eQTL SNP. SNPs are ordered left to right by their genomic position. Here, the two consensus SNPs (SNPs prioritized by at least two tools) are in near-perfect LD with the *SPNS1* eQTL lead SNP, suggesting that at this locus most likely alters PD risk through *SPNS1* gene expression.

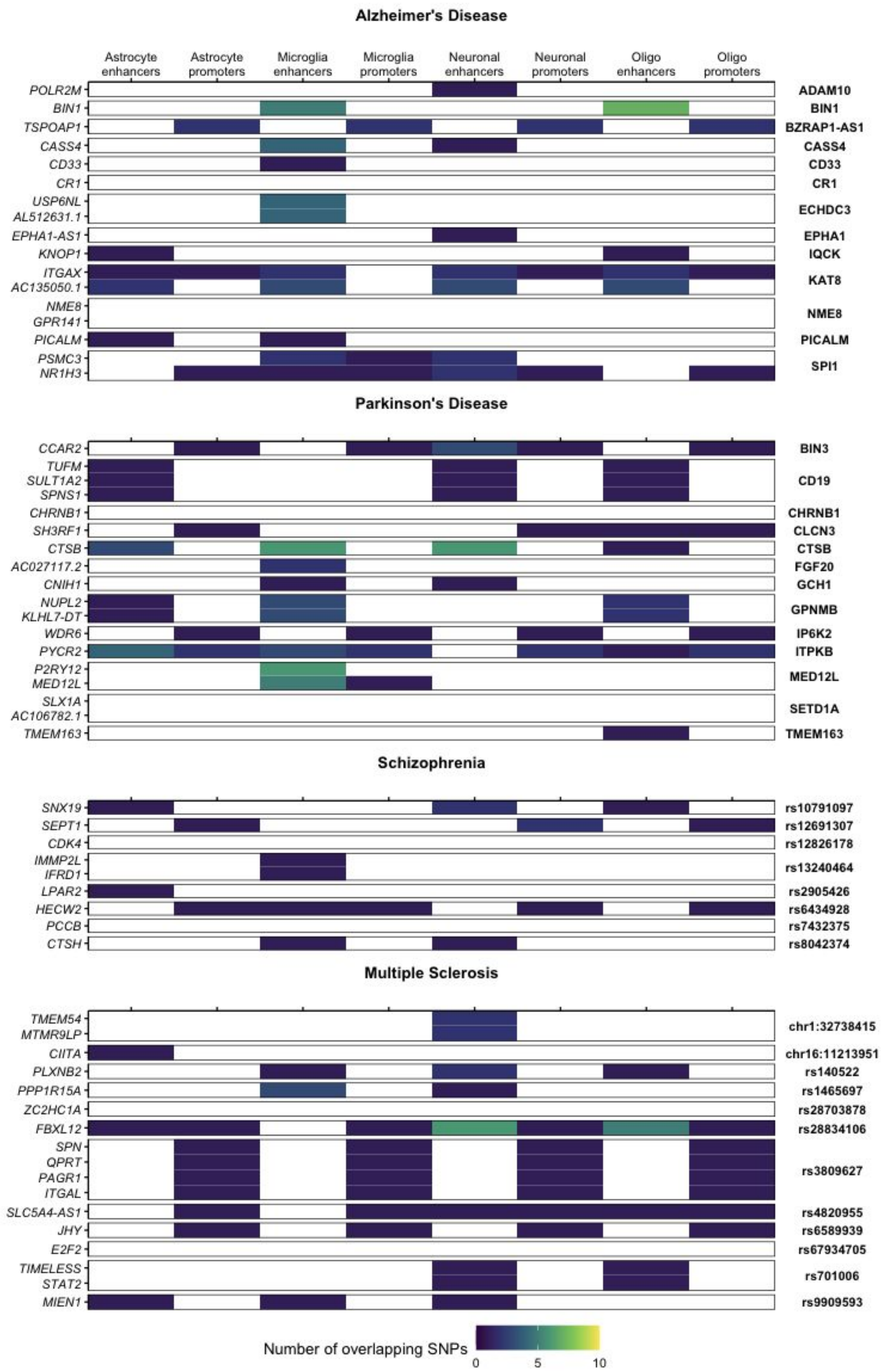


**Supplementary Figure 25: Overlap of colocalized microglia eQTLs with epigenomic features.**

Regions defined by Nott et al (2019) as cell-type specific promoters and enhancers were overlapped with SNP sets for each colocalizing microglia QTL - GWAS locus. SNP sets consisted of the lead GWAS SNP, the lead QTL SNP and any fine-mapped consensus or credible SNPs. Results are summarized here by the number of SNPs in the set that overlap with a particular feature type.
